# Nanotopography controls single-molecule mobility to determine overall cell fate

**DOI:** 10.1101/2020.07.23.191858

**Authors:** Marie FA Cutiongco, Paul M Reynolds, Christopher D Syme, Nikolaj Gadegaard

**Author notes:** These authors contributed equally to this work (Marie FA Cutiongco, Paul M Reynolds, Christopher D Syme).

## Abstract

The addition of nanoscale distortion to ordered nanotopographies consistently determines an osteogenic fate in stem cells. Although disordered and ordered nanopit arrays have identical surface areas, array symmetry has opposite effects on cell fate. We aimed to understand how cells sense disorder at the nanoscale. We observed effects in the early formation of cell and focal adhesions that controlled long-term cell fate. Disordered nanopits consistently yielded larger focal adhesions at a faster rate, prompting us to investigate this at the molecular scale. Super-resolution microscopy revealed that the nanopits did not act as nucleation points, as previously thought. Rather, nanopit arrays altered the plasma membrane and acted as barriers that changed molecular diffusion. The local areas corralled by four nanopits were the smallest structures that exerted diverging effects between ordered and disordered arrays. Heterogeneity in the local area on disordered arrays increased the proportion of fastest and slowest diffusing molecules. This resulted in higher quantity, more frequent formation and clustered arrangement of nascent adhesions, i.e., the modular units on which focal adhesions are built. This work presents a new pathway to exploit nanoscale sensing to dictate cell fate.

## Introduction

For over a decade and a half, precisely engineered nanopit arrays with different geometrical arrangements have been shown to elicit robust yet opposing biological effects. When tested on adult mesenchymal stem cells, nanopits in an ordered square array promote multipotency, while a subtle distortion in nanopit arrangement pushes cells toward an osteogenic lineage^1–15^. Changes in transcriptional and metabolic programmes across multiple cell lines underpin this phenotypic switch^2,3,5,8,15^. The cell and nuclear morphologies are also different between cells on ordered and disordered nanopit arrays. There is a recurring pattern in the effects of these nanopit arrays on cell adhesion and contractility, as well as on focal adhesion size. Specifically, disordered nanopit arrays lead to focal adhesion maturation, stress fibre formation and higher cell spreading, to induce the osteogenic fate. These three cornerstones highlight the development of intracellular tension as the differentiating factor between disordered and ordered nanopit arrays^6,14,15,17^.

The results of these studies are incongruous with the fact that both the ordered and disordered nanopit arrays are engineered to occupy the same total surface area. At the macroscopic level, cells should not be able to differentiate between the two types of nanopatterns. Meanwhile, the possibility of nanoscale effects triggered by nanopatterns has been suggested by prior studies, but remains poorly understood. Preliminary work using electron microscopy showed that nanopits in an ordered square array restricted cellular extensions, such as filopodia, to the spaces located between nanopits^8,17^. Previous reports also showed that cells on ordered nanopit arrays formed small focal adhesions that were limited to the spaces between nanopits^18,19^; however, these phenomena were measured using diffraction-limited techniques. These two studies support our working hypothesis regarding the manner in which nanopit arrays affect cell behaviour, i.e., that, compared with a flat surface, the nanopits serve as non-adhesive patterns in an otherwise adhesive area, and that the nanopit edges may act as nucleation sites that trigger integrin activation and clustering to enhance the formation of focal adhesions.

Our hypothesis mirrors the mechanism of constrained integrin clustering that arises from nanotopographies such as gratings^20^, tubes^21^ or pores^22^. Nanotopographies are suggested to exert cellular effects by effectively confining integrins within the maximum critical distance of 70 nm^23,24^. By controlling the availability, size and geometries of the areas available for integrin binding and clustering, nanotopographies affect mechanosensing and mechanotransduction via the focal adhesions^16^. The effect of the geometric variation of nanotopographies is less well known, but may occur analogously to distorted integrin ligands. Distortion in the order of integrin ligands spaced farther than 70 nm apart is postulated to yield variation in local inter-ligand spacing, with some distances falling well below the 70 nm threshold^25^. A similar mechanism may be at play on the nanopit arrays, with the disordered nanopit arrays providing a higher percentage of inter-pit distances below the 70 nm threshold. Therefore, with the lack of symmetry, a disordered nanopit array provides more directionalities for undisrupted focal adhesion growth compared with an ordered nanopit array.

In this study, we addressed how cells sense the nanoscale order of the nanopit arrays. We examined the effects of nanopit arrays via direct measurements at the nanoscale using techniques with a high spatio-temporal resolution. Our findings contradict long-held hypotheses regarding the mechanism of nanoscale sensing of nanopit order, which suggests the importance of nanopit edges as nucleation sites. In contrast, our results suggest that the quantity, spatial distribution and assembly of focal adhesion components are modulated by the nanopits and their arrangements on molecular mobility.

## Results

### Nanopits alter cell attachment, spreading, migration and differentiation

We fabricated substrates patterned with nanopits with a diameter of 120 nm, a depth of 100 nm and a nominal centre-to-centre spacing of 300 nm (Figure 1a). Nanopits were arranged in two different ways: in a square array (SQ) or displaced from the square array by a maximum of 50 nm in a so-called “near-square” arrangement (NSQ). Collectively, we shall refer to these arrangements as nanopatterns. An unpatterned (FLAT) substrate was included as a control. We used a pre-osteoblast cell line (MC3T3) in these experiments, as these cells completely replicate the previously observed osteogenic response of mesenchymal stem cells, osteoblasts and fibroblasts on FLAT, SQ and NSQ^10^ but exhibit the improved transfectability that is required for single-molecule imaging.

**Figure 1.**
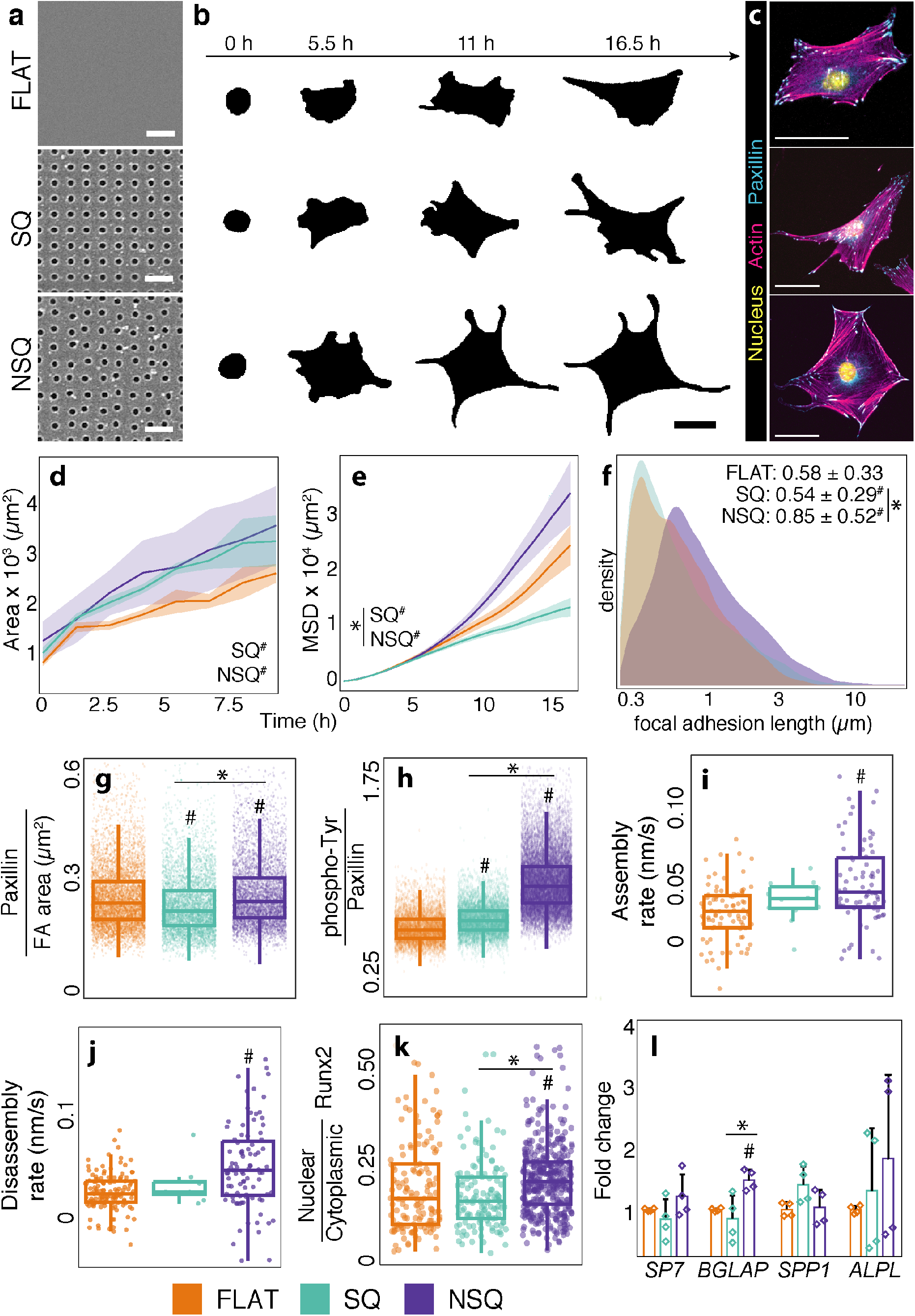
Nanopits dictate cell attachment, spreading, migration and differentiation. **a**, Arrays of nanopits with variations in geometry (“nanopatterns”) were used as culture substrates. An unpatterned (FLAT) substrate was used as a control. The nanopatterns consisted of pits with a diameter of 120 nm, a depth of 100 nm, a nominal spacing of 300 nm and arrangement in a square (SQ) array or displacement by up to 50 nm from the square array (NSQ). **b**, MC3T3 pre-osteoblast cells showed distinct attachment and spreading phenotypes on nanopatterns. Representative outlines of cells are shown. Scale bar, 5 μm. **c**, Fluorescence images of MC3T3 pre-osteoblast cells on nanopatterns. Nuclei are shown in yellow, actin filaments are shown in magenta and paxillin is shown in blue. Scale bar, 50 μm. **d**, The cell area on the nanopatterns varied over time. n = 16/11/11 for FLAT/SQ/NSQ in three independent experiments. **e**, Migration of cells on nanopatterns measured over time using mean squared displacement (MSD). n = 81/91/148 for FLAT/SQ/NSQ in four independent experiments. **f**, Distribution of focal adhesion length across multiple time points in response to nanopatterns. N = 7410/7227/9093 focal adhesions from 2/2/4 cells across two biological replicates. Focal adhesions were visualised in real time using cells transfected with GFP-FAK. Images were acquired at 5 min intervals for 35 min. **g–h**, The static focal adhesion properties changed in response to nanopatterns. n = 73/61/59 for (f), and n = 71/111/253 cells for (g) from FLAT/SQ/NSQ in two independent experiments. **i–j**, The dynamic properties of focal adhesions were altered by nanopatterns. n = 2/2/4 cells for FLAT/SQ/NSQ in two independent experiments. Focal adhesions were visualised in real time using cells transfected with GFP-FAK. Images were acquired at 5 min intervals for 35 min. **k–l**, Functional response of cells on nanopatterns in the absence of osteogenic biochemicals. Osteogenic differentiation was assessed by measuring the nuclear translocation of RUNX2 (n = 137/245/409 cells for FLAT/SQ/NSQ in one independent experiment) at day 2, and the expression of osteogenic genes at day 7 (n = 4 in two independent experiments). Genes are arranged in the order of increasing osteoblastic maturity, with *SP7* encoding the transcription factor osterix, *BGLAP* encoding osteocalcin, *SPP1* encoding osteopontin and *ALPL* encoding alkaline phosphatase. The line plots show the mean ± confidence interval, the boxplots show the median and interquartile range with minima and maxima at the whiskers, the dots indicate individual data points and summary statistics are given as the median ± median absolute deviation. # Denotes statistical significance against FLAT, and * denotes statistical significance between indicated pairs from one-way ANOVA with Tukey’s post hoc test. Exact *P* values are given in Supplementary Data 1.

The changes in morphology triggered by nanopits emerged rapidly. After seeding, pre-osteoblast cells on nanopatterns exhibited a profoundly changed cell morphology during spreading (Figure 1b–c, Supplementary Figure 1). Cells on FLAT exhibited the conventional adherent cell shape, with polarisation, whereas cells on nanopatterns had larger areas with extensive filopodia. Over a 10 h period, the cell area increased rapidly on both nanopatterns compared with FLAT (Figure 1d). Moreover, the effect of nanopatterns on cell migration was dependent on the arrangement of nanopits (Figure 1d). Compared with FLAT, the mean squared displacement (MSD) of cells was suppressed on the ordered SQ pattern but was accelerated on the disordered NSQ pattern.

The balance between cell adhesion and cell movement is tightly governed by focal adhesions. Here, we found that these adhesions were altered on all substrates. The length of the focal adhesions was increased on both nanopatterns compared with FLAT, with focal adhesions being longer on NSQ (Figure 1f). The intensity of paxillin (Figure 1g) and phosphorylated tyrosine (Figure 1h) staining within focal adhesions was also significantly increased on NSQ compared with SQ and FLAT. This ties in with our observations of focal adhesion dynamics, where focal adhesions on NSQ exhibited the highest average disassembly and assembly rates compared with FLAT (Figure 1i–j). In addition to indicating active mechanosensing and mechanotransduction, tyrosine phosphorylation in focal adhesion components drives for the formation of these structures, cell migration and cellular contraction^26–29^. The focal adhesion formation rates were statistically similar on FLAT and SQ.

These early changes in cell morphology translated into distinct long-term cell functionalities. Overall, NSQ robustly differentiated pre-osteoblast cells (even in the absence of biochemical induction). As early as day 2, the nuclear translocation of the osteogenic transcription factor Runx2 was significantly increased in NSQ compared with FLAT or SQ (Figure 1k). Osteogenesis was sustained by NSQ for 28 days, a time point at which the expression of the genes encoding for osteogenic markers osterix *(SP7),* osteocalcin *(BGLAP),* osteopontin *(SPP1)* and alkaline phosphatase *(ALPL)* was significantly enhanced compared with that observed for FLAT and SQ (Figure 1l).

Nanopatterns induced a systematic change in cell phenotype, which was apparent at multiple levels and time scales. The controlled distortion in nanopit arrangement that is present in NSQ stimulated cell spreading, rapid migration and focal adhesion formation and activation. The morphological changes induced by NSQ culminate in osteogenic differentiation in the absence of biochemical inducers. In contrast, we found no measurable change in cell function in the case of SQ.

### Nanopits are not nucleation sites for focal adhesions

It is well known that cells sense and respond to their microenvironment through focal adhesions. In turn, the transduction of external signals through focal adhesions alters cell behaviour. Our results highlight the correlation between focal adhesion characteristics and cell fate determination, especially regarding the osteogenic response induced by NSQ. Thus, our aim was to establish the effect of the presence of nanopits on focal adhesion assembly. In particular, we examined the long-held hypothesis that nanopits confine integrin activation to their edges, to initiate the formation of focal adhesions, similar to the observations regarding the effects of nanogratings or nanotubes^20–22^.

We used stochastic optical reconstruction microscopy (STORM), which enables the observation of the effects of nanopits on focal adhesions down to the nanometre scale. We used STORM to observe paxillin, which is one of the earliest adapter proteins that are recruited to nascent adhesions^30,31^ (Figure 2a). We focused our analysis on regions that were devoid of mature focal adhesions (>1 μm in length), which allowed the assessment of the onset of focal adhesion formation.

**Figure 2.**
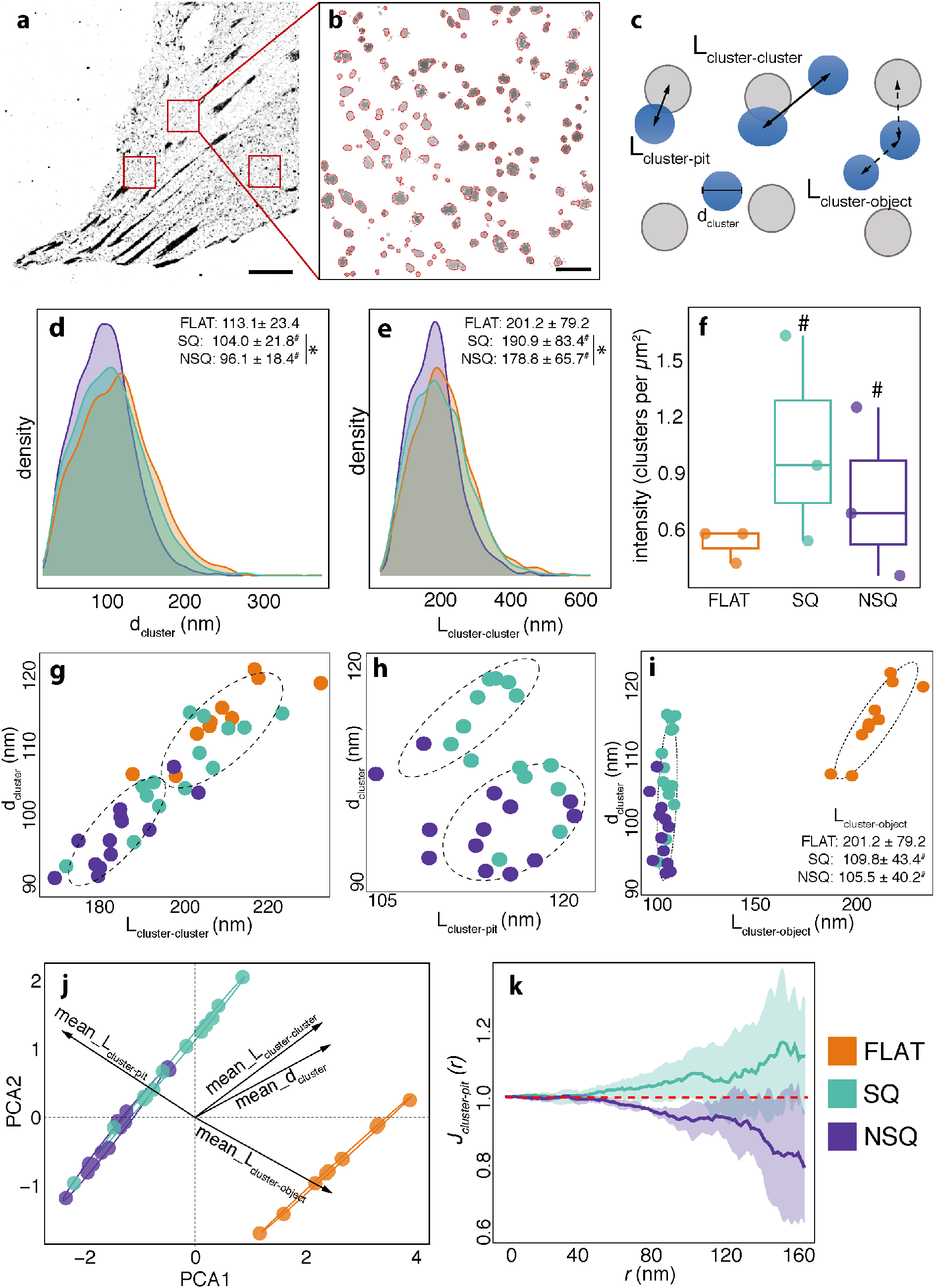
Nanopits alter the distribution and organisation of nascent adhesions. **a–b**, Paxillin was imaged using super-resolution microscopy. The red squares denote the exemplar regions (4 × 4 μm) in which paxillin molecules or clusters were analysed. Scale bar, 5 μm. The inset shows an exemplar region that was analysed for paxillin clusters (black outline). Scale bar, 500 nm. **c**, Diagram showing the interactions measured between paxillin clusters (in blue) and nanopits (in grey). d_cluster_ denotes the diameter of paxillin clusters, L_cluster–cluster_ denotes the distance between a paxillin cluster and the nearest cluster, L_cluster–pit_ denotes the distance between a paxillin cluster and the nearest nanopit, and L_cluster–object_ denotes the distance between a paxillin cluster and the nearest object, either a cluster or a pit. **d**, Distribution of d_cluster_ across different substrates. **e**, Distribution of L_cluster–cluster_ across different substrates. **f**, Intensity of paxillin clusters (counts per area) across different substrates. **g**, Scatterplot of d_cluster_ against L_cluster–cluster_ across different substrates. Paxillin clusters segregated into two different groups using an unsupervised k-means clustering algorithm. **h**, Distribution of L_cluster–pit_ across different nanopits. **i**, Scatterplot of d_cluster_ against L_cluster–object_ across different nanopits. Paxillin clusters segregated into two different groups using an unsupervised k-means clustering algorithm. **j**, Segregation of data points into groups using d_cluster_ and all nearest neighbour distances using a principal component analysis (PCA). PCA1 and PCA2 denote the first two dimensions obtained in the analysis. The contribution of each feature to the principal components are given in Supplementary Figure 7. **k**, Spatial dependence of paxillin clusters on nanopits. *J_cluster,pit_ (r)* is a summary of two measures: the probability of obtaining a distance *r* between a typical paxillin cluster and the nearest nanopit; and the probability of obtaining a distance *r* from any fixed location to the nearest pit. In particular, *J_cluster,pit_ (r)* > 1 indicates a regular pattern between clusters and pits; *J_cluster,pit_ (r)* < 1 denotes clustering between clusters and pits; and *J_cluster,pit_ (r)* = 1 denotes independence in the spatial pattern between paxillin clusters and pits. See the Materials and Methods for details. The dots indicate data points averaged from each analysed region. Summary statistics are given as the median ± median absolute deviation. Ellipses with dashed lines represent groupings of data calculated using unsupervised machine learning. Lines show the empirical *J(L_cluster,pit_),* with shaded areas denoting the confidence interval. n = 11/15/13 regions analysed for FLAT/SQ/NSQ obtained from three independent experiments; * denotes statistical significance between the indicated pairs from one-way ANOVA with Tukey’s post hoc test. Exact *P* values are given in Supplementary Data 1.

First, we addressed the hypothesis that nanopits affect focal adhesion formation by acting as nucleation sites. Under this proposition, two phenomena should emerge: 1) a higher number of paxillin molecules should be present on nanopit arrays compared with FLAT; and 2) paxillin molecules should be positioned near a nanopit. Our evidence opposed these expectations, as we observed that the global intensity of paxillin molecules (count per unit area) was comparable among all substrates (Supplementary Figure 2), with no preference for the nanopit edge. Furthermore, at the nanopit level, we found no relationship between paxillin molecule density and distance to the nanopit edge (Supplementary Figures 3 and 4, and Supplementary Note 1). Importantly, our results negated the hypothesis that the nanopits act as nucleation sites for the initial formation of focal adhesion components.

### Nanopits direct the size and spatial distribution of nascent adhesions

Since nucleation of focal adhesions at the nanopits is not responsible for the difference between SQ and NSQ, we wanted to understand the genesis of focal adhesions. To do so, we studied the effect of nanopits on the formation of nascent adhesions, which are the modular units of cell–matrix adhesions^32^. To examine this hypothesis, we changed our analytical approach to focus on the nascent adhesions, as defined by paxillin clusters, rather than individual molecules (Figure 2b). The analysis of paxillin clusters provides insights into the genesis of focal adhesions, as dictated by the nanopatterns. We identified individual clusters of paxillin using an unsupervised density-based clustering algorithm^33^ (see Materials and Methods for details). The paxillin cluster characteristics and spatial distribution were then measured to obtain information on the effect of nanopatterns on their spatial arrangement (Figure 2c). The diameter of paxillin clusters (d_cluster_) reflected the effect of nanopits on the size of nascent adhesions and inferred the amount of integrin activation. Although we observed a wide range (30–500 nm) of d_cluster_, the median value across all substrates was consistent with the reported size of the nascent adhesions (110 nm) initiated by liganded integrins^34^ (Figure 2d). The median paxillin cluster diameter was largest on FLAT and smallest on NSQ.

To measure the spatial distribution of paxillin clusters, we measured the pairwise distance between paxillin clusters and their nearest neighbouring cluster (L_cluster–cluster_). L_cluster–cluster_ provides information on the density of the packing and arrangement of paxillin clusters across different substrates. The L_cluster–cluster_ was longest on FLAT and shortest on NSQ, which indicated the tight packing of paxillin clusters on the latter pattern (Figure 2e). The large paxillin clusters spaced far apart observed on FLAT translated into the lowest intensity of paxillin clusters across all substrates (Figure 2f, Supplementary Figure 5). Both SQ and NSQ showed a higher intensity of paxillin clusters, indicating the formation of a greater number of nascent adhesions. On average, both FLAT and SQ had large paxillin clusters spaced farther apart compared with NSQ. An unsupervised clustering algorithm reflected the contrast in packing density between FLAT and NSQ using only d_cluster_ and L_cluster–cluster_ measurements (Figure 2g).

The distance between a paxillin cluster and its nearest neighbouring nanopit (L_cluster–pit_) was then measured to determine how nanopits alter paxillin cluster distribution. We found no significant difference in L_cluster–pit_ between SQ and NSQ (Figure 2h). For comparison with FLAT, we extended the L_cluster–pit_ measurement to include the distance from a paxillin cluster to its nearest neighbouring object (either another paxillin cluster or a nanopit, L_cluster–object_); i.e., for FLAT, L_cluster–cluster_ is the same measure as L_cluster–object_. The L_cluster–object_ value allowed us to gauge the effect of adding nanopits to the available empty space (Supplementary Figure 6). On both nanopit arrays, the L_cluster–object_ was significantly smaller than was the L_cluster–object_ on FLAT. This means that the addition of nanopits, regardless of arrangement, led to a greater compression of paxillin clusters, with less empty space separating them (Figure 2i).

The collective changes in d_cluster_, L_cluster–cluster_, L_cluster–pit_ and L_cluster–object_ induced by the different substrates were assessed using principal component analysis (PCA). PCA discriminated between FLAT, SQ and NSQ (Figure 2j, Supplementary Figure 7). FLAT was clearly separable because of its stark difference in L_cluster–object_ compared with SQ and NSQ. SQ was separated from NSQ by the larger d_cluster_ and longer L_cluster–cluster_. As measured based on paxillin clusters, our results demonstrated how nanopits form a greater number of nascent adhesions with smaller sizes. Importantly, Nanopits constrain the free space of the substrate to increase the density of the nascent adhesions compared with FLAT, which is a mechanism through which the formation of focal adhesions can be accelerated.

### Nanopit arrangement dictates the spatial pattern of nascent adhesions

We used robust spatial statistics^35^ to study systematically the large-scale spatial patterns of paxillin clusters on FLAT, SQ and NSQ. We first confirmed that the intensity and correlation of paxillin clusters exhibited a spatial pattern (i.e., a non-random pattern) (Supplementary Table 1). Further analysis of correlation and spacing using summary statistics revealed similarities between FLAT, SQ and NSQ (Supplementary Figure 8). Regardless of the substrate, the spatial patterns in paxillin clusters were similar regarding correlation (the average number of paxillin clusters found within a certain distance from a typical paxillin cluster) and spacing (the number of nearest neighbours within a certain distance from a typical paxillin cluster).

We then used spatial statistics to determine if the spatial distribution of paxillin clusters was dependent on the spatial distribution of nanopits. The intensity of paxillin clusters showed no dependence on SQ or NSQ nanopit locations (Supplementary Table 2). Conversely, the correlation and spacing of paxillin clusters and nanopits were significantly different from an independent and random spatial pattern, which confirmed spatial dependency (Supplementary Figure 9). In particular, we observed that the spacing between paxillin clusters and pits was significantly different on SQ vs. NSQ. In particular, the *J_cluster–pit_ (r)* function showed that the order of nanopits affected the spatial distribution of paxillin clusters. *J_cluster–pit_ (r)* is a dimensionless ratio of two probabilities: (1) the probability that the distance between a cluster and the nearest nanopit is greater than a distance *r*; and (2) the probability of finding a nanopit within a distance *r* from any arbitrary location. At a particular distance *r*, the *J(r)* function provides information on the prevalence of nearest neighbours compared with empty distances. When *J_cluster–pit_ (r)* > 1, clusters are spaced regularly from nanopits at distances ≤ *r*. When *J(r)* < 1, clusters and nanopits are spatially aggregated at distances ≤ *r*. When *J(r)* = 1, there is no dependence of cluster arrangement on nanopit location (see Materials and Methods for a detailed explanation).

The *J_cluster–pit_ (r)* function between clusters and nanopits (*J_cluster–pit_ (r)*) exhibited a clear divergence between NSQ and SQ (Figure 2k). NSQ showed *J_cluster–pit_ (r)* < 1, suggesting an association between paxillin clusters and nanopits and spatial clustering between these objects. On SQ, the *J_cluster–pit_ (r)* > 1 denoted repulsion between paxillin clusters and nanopits and a regularity in the arrangement of these objects. The overlay of nanopits in a square array on paxillin clusters from FLAT failed to replicate the spatial arrangement of paxillin clusters on SQ (Supplementary Figure 10)

Our systematic analysis pointed to the effect of nanopatterns in controlling the density and distribution of components needed for focal adhesion assembly and not by providing nucleating sites. The nanopits by themselves are associated with enhanced formation of nascent adhesions, which are closer packed than on a flat substrate. Additionally, we found that SQ disperses the clusters whereas NSQ leads to aggregation of clusters. Therefore, the order of nanopit arrangement dictates this packing into an irregular or regular pattern.

### Variation of local areas bound by nanopits permits the localisation of more nascent adhesions

The data obtained thus far have shown that (1) nanopits affect nascent adhesion properties and spatial arrangement relative to FLAT; and (2) the subtlety of nanopit arrangement dictates the spatial arrangement of nascent adhesions. Although we established how nanopits differ from FLAT, it the discernible characteristics of SQ and NSQ that led to opposite spatial arrangements remained less clear.

To address this question, we examined STORM reconstructions of paxillin in correlation with the electron microscopy observations of the nanopits (Figure 3a–d). Using this visualisation, two trends were apparent. First, the local areas on SQ and NSQ varied significantly. The smallest local area definable on the nanopatterns was the enclosure with four nanopits at its vertices, which we termed the bounding box (Figure 3e). The area of bounding boxes on SQ was constant at 300 × 300 nm = 9 × 10^4^ nm^2^. Although the average bounding box area on NSQ was equivalent to that of SQ, NSQ boxes ranged in area from 200 × 200 nm = 4 × 10^4^ nm^2^ to 400 × 400 nm = 16 × 10^4^ nm^2^.

**Figure 3.**
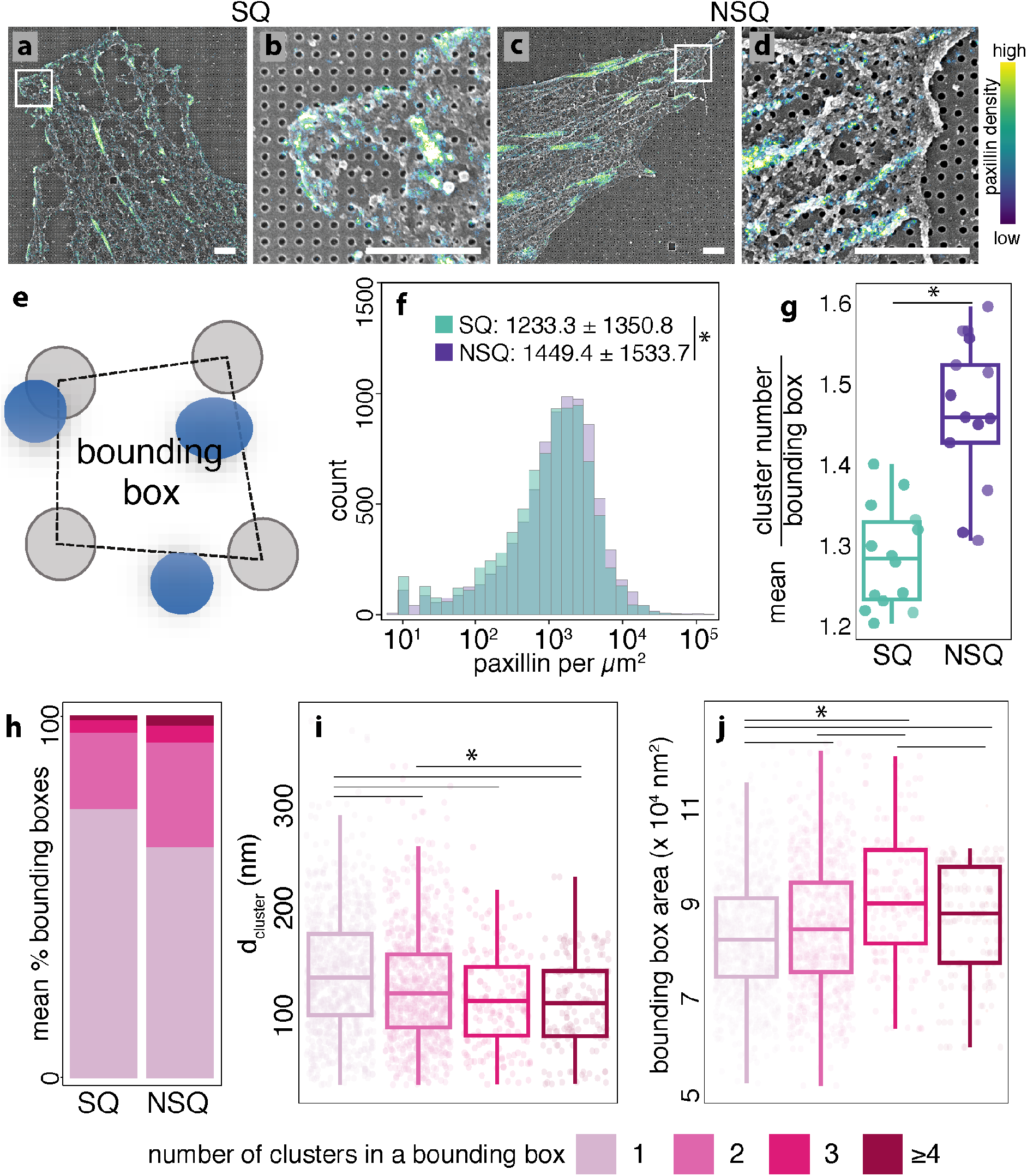
The variability in NSQ bounding box area induces higher paxillin cluster formation. **a–d**, Representative images of paxillin (visualised using super-resolution microscopy) and nanopits (electron microscopy). The white boxes indicate the magnified regions in the bottom panel. Scale bar, 2 μm. **e**, A bounding box defined by four nanopits at its vertices. The bounding box areas for SQ were constant at 9 × 10^4^ nm^2^. The bounding box areas for NSQ ranged from 4 to 16 × 10^4^ nm^2^. Nanopits are shown in grey, paxillin clusters are shown in blue and the bounding box is indicated by a dashed line. **f**, Density of paxillin localisations in bounding boxes. The data are presented as the number of paxillin molecules in a bounding box, normalised to that bounding box area. For visualisation, the x-axis is presented in the log scale. **g**, Average number of paxillin clusters found in a bounding box. **h**, Proportion of bounding boxes containing specific amounts of paxillin clusters. The data are shown as a percentage of the total number of bounding boxes analysed. The colour scheme indicates the number of clusters found in the bounding box. **i**, Distribution of NSQ bounding box area filled with different amounts of pre-FA clusters. The colour scheme denotes the number of clusters found in the bounding box. **j**, Distribution of the diameters of clusters found within bounding boxes filled with different amounts of pre-FA clusters. The colour scheme denotes the number of clusters found in the bounding box. The bar plots show the mean ± standard deviation, with individual points denoting mean values from individual regions. The boxplots show the median and interquartile range with minima and maxima at the whiskers, the dots indicate data points averaged from individual regions and summary statistics are given as the median ± median absolute deviation. n = 15/13 regions analysed for SQ/NSQ obtained from three independent experiments; * denotes statistical significance between the indicated pairs from ANOVA with Tukey’s post hoc test. Exact *P* values are given in Supplementary Data 1.

Second, correlative microscopy revealed that the number of paxillin clusters varied locally. We applied the bounding boxes to assess the effects of SQ and NSQ on the local density of paxillin clusters (Figure 3f). On average, we observed more paxillin molecules per μm^2^ on NSQ vs. SQ. This translated into a higher average count of paxillin clusters in each bounding box on NSQ compared with SQ (Figure 3g). These results were unique to the nanopatterns, as they could not be replicated by overlaying a nanopit pattern with a square arrangement over the paxillin clusters obtained from FLAT (Supplementary Figure 10). A comparison of the nanopatterns revealed that NSQ had the highest average proportion of bounding boxes with multiple paxillin clusters (Figure 3h). We postulate that the variation in the bounding box area on NSQ expands the local area within which paxillin clusters can reside. In fact, this was supported by a decrease in the average d_cluster_ within bounding boxes with more paxillin clusters (Figure 3i). Furthermore, we observed that a significant increase in the average bounding box area was accompanied by a higher number of paxillin clusters (Figure 3j).

We established here how the variation in the bounding box area stemming from the distortion of nanopit arrangement on NSQ increased the number of paxillin clusters residing within the local vicinity. On SQ, in contrast, the constant bounding box area provided no opportunities to accommodate higher amounts of paxillin clusters.

### Nanopits suppress molecular mobility

The results so far shows that, regardless of arrangement, nanopits change the number, size and spatial distribution of paxillin clusters. However, the variation in the bounding box areas of NSQ was the crucial parameter in determining the co-localisation of multiple paxillin clusters within the same local area. These results have important implications for understanding the manner in which focal adhesions are formed. These findings, together with the nanopattern-induced effects on focal adhesion dynamics and size (Figure 1), led us to explore dynamics at the nanometre length scale. We examined how nanopits affect the spatio-temporal characteristics of paxillin and its consequences on focal adhesion assembly using single-particle tracking. We also tracked the movement of the transferrin receptor (TfR), a transmembrane protein, to infer the adhesion-independent effects of nanopatterns on the plasma membrane.

For both paxillin and TfR, we observed clear differences in the characteristics of molecular tracks depending on nanopatterns (Supplementary Figures 11 and 12). Across all nanopatterns, paxillin and TfR track velocity was maximised, whereas confinement was minimised on FLAT. The paxillin and TfR tracks on NSQ were faster, less confined and more tortuous than those observed on SQ. Furthermore, overlaying tracks with nanopits revealed that the movement of TfR and paxillin was pinned to nanopit edges and hopped between nanopits (Figure 4a). Similar to paxillin clustering, we observed that nanopits dictated the spatial positioning of single molecules. Using ensemble averages of track MSD, we classified the movement of paxillin and TfR under the control of the nanopatterns. The diffusion state is typically inferred by estimating the α value using the following equation, which shows the dependency of MSD on time:

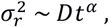

where 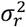 is the time-averaged ensemble MSD, *D* is the diffusion coefficient and *t* is the elapsed time. On FLAT, both paxillin and TfR showed normal Brownian motion, as defined by α = 1, where α indicates the dependence of MSD on time (Figure 4b–c). Conversely, both nanopatterns induced anomalous diffusion (α < 1) of paxillin and TfR. The largest suppression of paxillin and TfR movement was observed on SQ, in agreement with macroscopic data on focal adhesion dynamics (Figure 1i–1j). Nanopits, regardless of arrangement, acted as immobile obstacles to curtail free diffusion.

**Figure 4.**
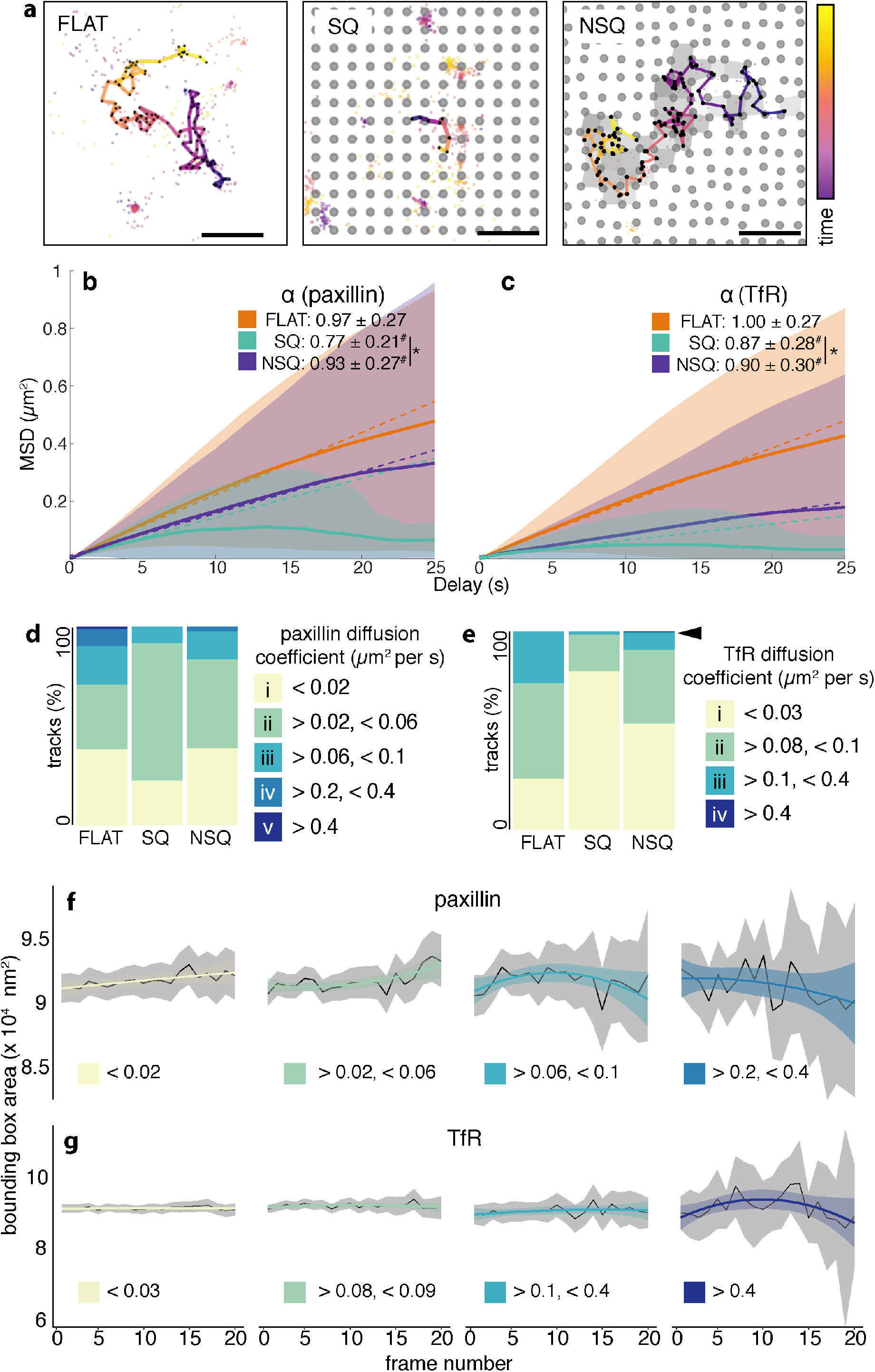
Disorder in nanopit arrangement changes the density and speed of diffusion of paxillin. **a**, Representative tracks of transferrin receptor (TfR) movement on FLAT and the nanopatterns. Mobile TfR and immobile TfR are shown as individual dots, nanopits are shown in grey circles and bounding boxes on NSQ are shown as grey polygons. The colour scale indicates the relative time within each diagram. **b–c**, Anomalous diffusion of paxillin and TfR induced by nanopatterns. α denotes the relationship between the time delay and the mean squared displacement (MSD) and determines the diffusive regime: α = 1 denotes normal diffusion, α > 1 denotes active or super-diffusion and α < 1 denotes anomalous or sub-diffusion. α values fitted from tracks are shown in the text. The solid line indicates the average of the ensemble MSD mean over time ± weighted standard deviation, and the dashed line shows the linear fit from which the estimated diffusion coefficient was obtained. # Denotes statistical significance against normal diffusion (α = 1), as tested using a two-tailed *t*-test, and against FLAT, as tested using one-way ANOVA with Tukey’s post hoc test. * Denotes statistical significance between the indicated pairs. Summary statistics are given as the mean ± standard deviation. **d–e**, Proportion of tracks within each diffusive subspecies modelled for paxillin and TfR. The colour scheme sows the different diffusive subspecies, each with its range of diffusion coefficient. The black arrow indicates the presence of the fastest TfR subspecies on NSQ. The molecular subspecies with increasing diffusion coefficients are indicated by the categories (i), (ii), (iii), (iv) and (v). The statistical analysis used to compare the proportion of tracks in each subspecies is provided in Supplementary Data 1. **f–g**, Correlation between the diffusion coefficient of paxillin or TfR and the bounding box area on NSQ. The black lines with a grey shaded area indicate the mean ± confidence interval from empirical data. The coloured lines and shaded regions denote a parabolic curve fitted to the dataset. Diffusive subspecies with increasing diffusion coefficients are indicated (i), (ii), (iii), (iv) and (v). For the whole figure, n = 864/1534/1409 tracks for FLAT/SQ/NSQ from one independent experiment for paxillin; and n = 2002/1031/1309 tracks for FLAT/SQ/NSQ from two independent experiments for TfR. # Denotes statistical significance against FLAT, and * denotes statistical significance between the indicated pairs from one-way ANOVA with Tukey’s post hoc test. Exact *P* values are given in Supplementary Data 1.

### Distortion of nanopattern arrangement augments the proportion of extreme diffusive states

In addition to estimating the diffusion regime, single-particle tracking can yield information about kinetics. This is useful for extrapolating the behaviour of paxillin, the movement of which is dominated by binding equilibria. The paxillin binding kinetics changes in response to distance from a focal adhesion^36^ and on the properties (orientation^37^ or dynamic state^38^) of the focal adhesion in which it resides. Measuring changes in the apparent diffusion coefficient, which was used here to categorise track movement into diffusion states, is an approach to determining changes in binding kinetics.^39^

Our findings regarding the range of diffusion subspecies for paxillin match those reported previously^38^ (Figure 4d; the full list is given in Supplementary Table 4). Paxillin dynamics showed less bias in distribution among subspecies i–iii on FLAT compared with all nanopatterns (Supplementary Data 1). FLAT and NSQ exhibited the highest number of slowest-moving paxillin (subspecies i), which was indicative of the tight association between the paxillin molecules that are normally found inside stable focal adhesions. Compared with SQ, NSQ exhibited an increased proportion of paxillin with higher diffusion coefficients (subspecies iii–iv). More paxillin molecules diffusing at a faster rate on NSQ than SQ imply more paxillin found in growing focal adhesions^38^. This increased proportion of rapidly moving paxillin is freely available, but transiently bound to focal adhesions, possibly representing a pool for easy recruitment and binding into a focal adhesion when required. The presence of faster-diffusing paxillin ties in with the rapidity of focal adhesion assembly and increased focal adhesion size observed on NSQ (Figure 1i & f). These results underpin the different rates of focal adhesion dynamics and cell migration on the nanopatterns.

A similar analysis of TfR diffusion states was performed (Figure 4e; the full list is given in Supplementary Table 2). TfR subspecies i–iii were more equitably distributed on FLAT compared with the two other nanopatterns (Supplementary Data 1). The largest proportion of slowest-moving TfR (subspecies i) was found on SQ. Similar to our observations regarding paxillin, NSQ induced an increasing proportion of TfR to move faster (subspecies ii–iii). The emergence of a new TfR subspecies (iv, black arrow) on NSQ, but not on SQ, is noteworthy and emphasised the effect of a disordered nanopit arrangement on increasing the number of faster-moving molecules. Overall, we showed that, for both paxillin and TfR, nanopatterns alter the distribution of trajectories to increase the likelihood of obtaining extremely slow and extremely fast molecules.

### The diffusion speed depends on the bounding box area

Our results on TfR movement argue in favour of the direct restructuring of the membrane triggered by nanopatterns. Together with the bounding box analysis presented in Figure 3, the emerging mechanism of nanopattern-induced effects coincided with the picket and fence model of membrane structure. In this model, the membrane is compartmentalised through transmembrane proteins that act as pickets and an actin-based membrane skeleton that acts as a fence.^40–42^ The picket and fence model was proposed as a means to explain the confined and hopping diffusion of membrane proteins. Based on this paradigm, we speculate that nanopits alter the plasma membrane to create compartments that mirror the bounding boxes. Therefore, we used the same bounding box analysis first shown in Figure 2 to correlate membrane structure and molecular mobility with NSQ bounding box variation (Figure 4f–g). The two slowest diffusing paxillin (subspecies i–ii) showed a slight increase in average bounding box area over time. This relationship was reversed for the fastest-diffusing paxillin (subspecies iii–iv), which exhibited a decrease in the average bounding box area over time. Moreover, most TfR subspecies (i–iii) showed no change in average bounding box area over time. Only the fastest TfR subspecies (v) showed a dependence on the decrease in the bounding box area over time. Our results revealed that the disordered arrangement of nanopits provides a variation in the local available area that redistributes diffusion species toward faster-moving molecules.

## Discussion

When considered at a scale similar to the size of a cell, both SQ and NSQ exhibit the same total area outside nanopits. However, multiple reports have demonstrated the effect of these nanopatterns at the microscale, as manifested in focal adhesions, the nucleus and the actin cytoskeleton. Here, we aimed to understand how cells sense nanoscale order. We used tools with high spatio-temporal resolution to interrogate the previously unexplored effects of nanopatterns at a comparable nanometre length scale. Throughout this study, a bottom-up mechanism for nanosensing emerged: nanopits and their arrangement altered focal adhesion construction, starting from their effects on single-molecule movement.

Our study supports a model in which nanopatterns direct the structure of the plasma membrane in a manner analogous to a picket and fence structure^40–42^. We postulate that the nanopits serve as stable pickets, which appear to repel molecules, whereas the fences (formally originating from a membrane-associated skeleton) delineate the membrane into the inter-picket space, which is analogous to our bounding boxes. The inter-picket space can vary in size from 30 to 230 nm^43^, which coincides with the possible size of bounding boxes on NSQ. Our results regarding sub-diffusive movement and our analysis of bounding boxes support this theory. The bounding box compartments in the plasma membrane sterically hinder molecular mobility and leads to a sub-diffusive regime^44^. Further suppression of free diffusion in the inter-picket space is afforded by a viscous plasma membrane, which is caused by the local interaction between transmembrane protein pickets and lipids. Within the bounding box, molecular confinement is also enhanced and, thus, the rate of molecular collision is increased^45^ and leads to enhanced aggregation and binding of focal adhesion components, such as integrins. The enhanced formation of nascent adhesions (measured from paxillin clusters) observed on the nanopatterns vs. FLAT supports this notion. Thus, nanopits restructure the plasma membrane to suppress diffusive regimes, induce different molecular states and promote molecular confinement. We speculate that the imprinting of the plasma membrane by the nanopatterns causes changes in molecular and cell behaviour. Previous work on artificial lipid bilayers showed that this is possible on gratings with a 360 nm width and a 320 nm depth^46^, while experiments confirmed the conformation of membranes into pillar structures^47^ that yield changes in lipid tension and membrane diffusion^48^. Further work aimed at dissecting this possibility would complement the results presented here.

Nevertheless, it appears that another phenomenon is in place to differentiate the mechanisms of formation of nascent adhesions based on NSQ bounding boxes. In contrast with SQ, which has a constant bounding box area, smaller NSQ bounding boxes were correlated with faster molecular diffusivity and fewer but larger nascent adhesions. The thermodynamic consequences of molecular crowding may be more prominent in smaller bounding boxes than in larger ones. The decrease in bounding box area triggers a concomitant increase in the free energy of the system that is required to confine molecules^49^. Consequently, the system will need to compensate, and it does so entropically by enhancing the interaction between molecules with *de novo* capacity for association (integrin heterodimerisation, for instance)^50^. The effect of a crowded membrane is less pronounced on larger bounding boxes. Moreover, together with slower-moving molecules (or longer residence times), this indicates a more careful nanoscale sensing and creates increased opportunities for the formation of multiple sites of nascent adhesions^51^.

Enhanced integrin clustering has consequences on mechanosignalling, which have been reported to occur more prominently on NSQ than SQ^3,10^. Increased integrin clustering was recently found to linearly increase its association with, and phosphorylation of, the critical effector protein focal adhesion kinase^52^. The existence of bounding boxes that are smaller than average induces an enhancement in molecular confinement and a consequent increase in mechanosignalling on NSQ, consistent with previous reports of higher levels of focal adhesion kinase phosphorylation on NSQ compared with SQ^3,10^. The phosphorylation of focal adhesion kinase is a critical event in the regulation of *in vitro* osteogenesis^53,54^, which is a consistent outcome of cell growth on NSQ (reported here and by others^2,4,6,10^). In addition to the formation of nascent adhesions, the variation in the bounding boxes that are present in NSQ and lacking in SQ crucially affects mechanosignalling and, ultimately, the cell phenotype.

The formation of multiple nascent adhesions in a bounding box enabled by the variations in the areas of NSQ translates into focal adhesion growth and maturation. While the formation of nascent adhesions primarily requires liganded integrins, its growth and stability require traction force^55,56^. Recent studies uncovered that this traction force is dependent not only on the distance between integrin ligands (maximum of 70 nm), but also on dimensionality^34^. A high density of integrin ligands arranged in isolated lines with a width of 30 nm spaced 250 nm apart yielded only transient nascent adhesions that fail to recruit paxillin, activate downstream signalling via pFAK or form actin stress fibres. Conversely, pairs of integrin ligand lines (distance of 80 nm) spaced 500 nm apart induced cell spreading, focal adhesion formation and downstream signalling. Despite the similar global ligand density, the availability of nascent adhesions in at least two dimensions is needed to allow stabilisation and linkage to the actomyosin machinery by the talin dimer. It appears that a similar phenomenon occurs on nanopit arrays, in which both the distance and dimensionality requirements for focal adhesion maturation are met more frequently on NSQ. Compared with SQ, clusters on NSQ were found to be closest to each other. The increased permissivity for forming more nascent adhesions within a single bounding box provides another opportunity for the distance criterion to be met. Moreover, the likelihood of the maturation of focal adhesions is lower on SQ than it is on NSQ because of the larger distances between nascent adhesions and the lower frequency of obtaining multiple nascent adhesions in one bounding box. In terms of dimensionality, as the spatial arrangement of nascent adhesions is opposite on SQ and NSQ. Intuitively, a disorganised or clustered arrangement of nascent adhesions on NSQ imposes less restrictions on the directionality of the bridging of another nascent adhesion for maturation into a focal adhesion. Conversely, the more regular array of nascent adhesions observed on SQ implies a maximum of two possible directions for bridging between nascent adhesions. In addition to increasing length, this mechanism underpins the stark enhancement in phosphorylated tyrosine levels and active signalling of focal adhesions on NSQ. Here we unravelled the sensing of order that occurs at the nanoscale (Figure 5). We established that cells respond to nanopits by modifying their molecular behaviour to modulate the manner in which adhesions are built, rather than by rigidly defining locations for the formation of adhesions (as was previously thought). Furthermore, we showed that the nanoscale sensing of order occurs at the smallest structure that can be varied between ordered and disordered nanopit arrangements (as defined by four nanopits), rather than at the single-nanopit level. The fundamental nature of the nanoscale sensing mechanism presented here, which occurs at the single-molecule level, broadens our understanding of how nanotopographies define cell fate.

**Figure 5.**
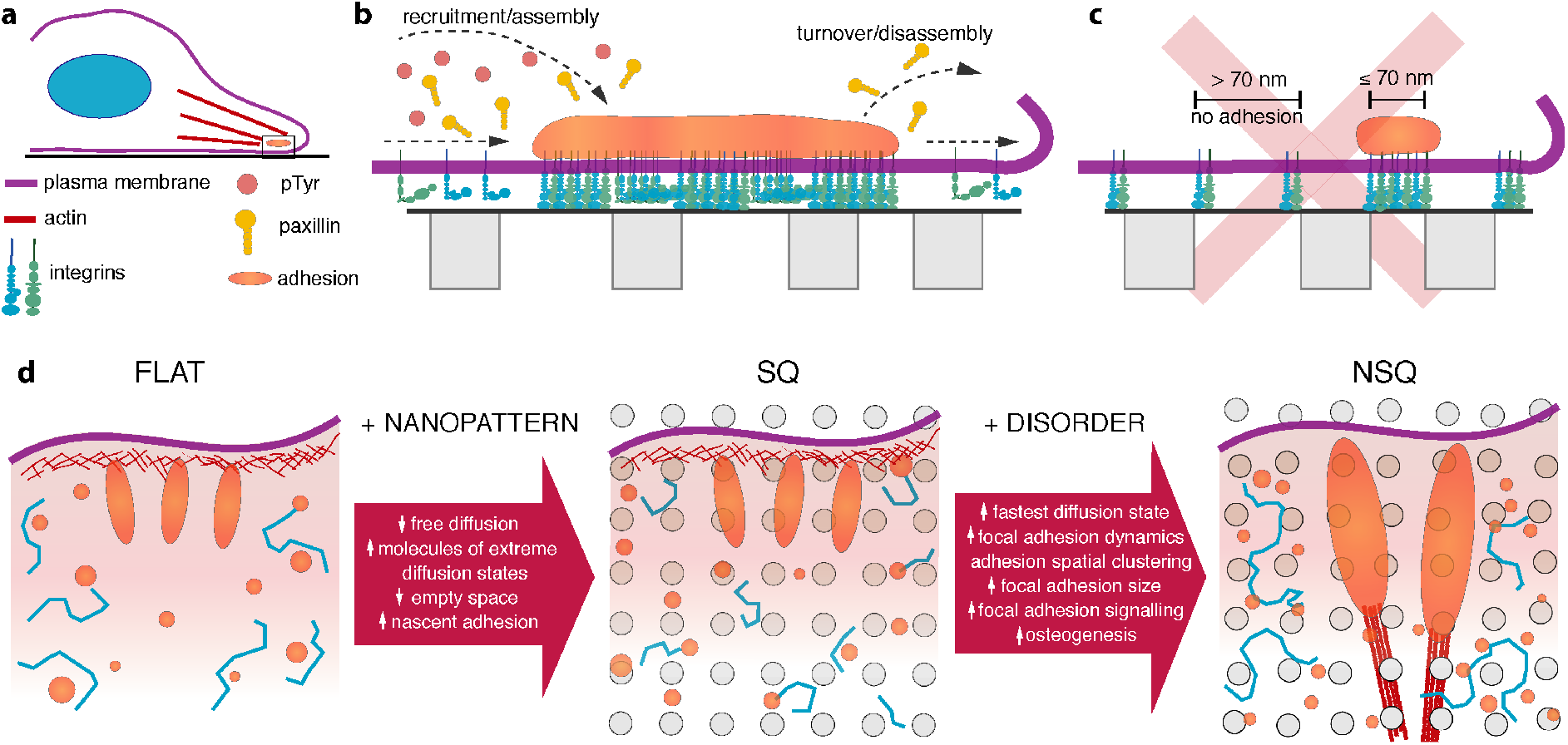
Nanoscale sensing of order. **a**, The effects of nanopatterns are first observed on focal adhesions, which exhibit changes in size, dynamics and signalling activity. **b**, The effects of nanopits emerge from changes in the diffusion of components into and out of nascent adhesions, which are the modular units of focal adhesions. **c**, Our study negates the hypothesis of previous studies, which suggests the emergence of focal adhesion formation from the nucleating effect of nanopits and the growth of adhesions between nanopits with distances constrained to 70 nm or less. **d**, The nanopits, regardless of order, suppress molecular diffusion by acting as obstacles to the free movement of molecules. A slower molecular movement provides more opportunities to form nascent adhesions. The addition of disorder in nanopit arrangement provides variability in bounding box area. In turn, the heterogeneity in the local area increases the number of fastest-diffusing states and the accommodation of multiple nascent adhesions. These changes propagate and render focal adhesions larger, more dynamic and more active at signalling, which culminates at osteogenesis on disordered nanopit arrays.

## Methods

### Nanofabrication of patterned coverslips

Glass coverslips (Menzel Glazer 1.5H) were solvent cleaned in an ultrasonic bath for 5 min each in acetone, methanol, isopropyl alcohol and water before drying and dehydration in a 180°C oven for 60 min. Polymethyl methacrylate (PMMA) resist (Elvacite 950K) was spun to a thickness of 100 nm, followed by a soft bake on a hotplate at 180°C for 2 min. A 10 nm aluminium charge conduction layer was evaporated before exposure of the nanodot patterns using a Vistec VB6 tool. The approximate write time for each coverslip was 30 min. After exposure, the charge conduction layer was removed by a 60 s dip in 2.6% tetramethylammonium hydroxide, followed by a 60 s RO water rinse. The exposed pattern was developed in a 2.5:1 mixture of methyl-isobutyl-ketone:isopropanol for 30 s at 23°C, followed by a 30 s rinse in isopropanol.

As PMMA is not compatible with the dehydration process needed for scanning electron microscopy (SEM), we fabricated alternative samples for correlative electron microscopy (CLEM). A 200 nm HSQ (Dow Corning) layer was spin coated onto clean coverslips and the film was annealed at 450°C for 10 min under nitrogen purge. Higher temperatures would improve the resulting silica film; however, the thin coverslips warped at temperatures above 500°C. A 200 nm layer of CSAR resist (Allresist) was spin coated onto the coverslips and soft baked at 150°C for 3 min. A 10 nm aluminium charge conduction layer was evaporated and the nanopatterns were exposed as described above. After exposure, the aluminium layer was removed and the nanopatterns were developed in amyl acetate for 30 s at 23°C, followed by a 5 s dip in o-xylene and two 30 s rinses in isopropanol. Any residual resist was removed by short RIE oxygen plasma at 100 W for 20 s. The exposed pattern was transferred into the cured HSQ layer via RIE using CHF_3_/Ar chemistry with an etch rate of approximately 30 nm/s. This followed the oxygen plasma and was run for 4.5 min, yielding an etch depth of approximately 130 nm. The CSAR resist was removed by overnight soak in SVC-14 at 50°C, followed by hot sonication for 10 min. Before cell culture, the coverslips were sterilised under UV for 20 min.

### Cell culture

Mouse pre-osteoblast cells (MC3T3, ATCC, passages 11–14) were cultured in alpha minimum essential medium (αMEM) with 10% foetal bovine serum, 100 U penicillin and 10 μg/ml streptomycin, then grown under the standard conditions of 37°C in a 5% CO_2_ environment.

### Cell functionality measures

Cells were allowed to differentiate in response to the nanopatterns without biochemical inducers of osteogenesis. Polystyrene patterned with the same nanopit arrays was used to study differentiation, to ensure the isolated effects of each nanopattern without confounding paracrine effects^57^. Cells were seeded onto patterned polystyrene at 5000 cells/cm^2^ and allowed to grow for either 2 days or 28 days. At 2 days of culture, cells were fixed with 4% paraformaldehyde then immunostained for Runx2 (Abcam ab76956, 1:250), followed by incubation with a fluorescently conjugated secondary antibody (ThermoScientific, 1:500), rhodamine phalloidin (ThermoScientific, 1:50) and DAPI (ThermoScientific, 1:5000). Patterned polystyrene samples were then mounted onto 0.17 μm thick coverslips prior to imaging. Images of cells were obtained using a 40× objective (numerical aperture (NA), 1.3). Single-cell profiling was performed with CellProfiler^58^ (v2.4, Broad Institute) using actin-stained images to delineate individual cells.

Differentiation was also assessed through the measurement of gene expression after 28 days of culture. Total RNA was obtained using the ReliaPrep RNA System (Promega) according to the manufacturer’s instructions. The expression of the osteogenic genes *SP7*, *BGLAP*, *SPP1* and *ALPL* was determined using 5 ng of total RNA and a one-step SYBR-based quantitative polymerase chain reaction kit (PrimerDesign). Gene expression assays were run on a BioRad CFX96 platform. Relative gene expression was first normalised to the levels of the 18S reference gene (Δ*C*_*T*_) and subsequently normalised to the gene expression levels detected on FLAT(ΔΔ*C*_*T*_). Fold changes are reported here as 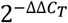. All primer sequences (Supplementary Table 5) were validated to have a single melt curve peak and a single amplicon with the expected size.

### Image-based cell profiling

Cells were grown on nanopatterns for 2 days. Subsequently, cells were fixed with 4% paraformaldehyde in phosphate-buffered saline (PBS), permeabilised then immunostained with the following antibodies: anti-Runx2 (Abcam ab76956, 1:250), anti-YAP (Cell Signalling Technologies 4912, 1:70), anti-phosphorylated Tyrosine (Cell Signalling Technologies 9415, 1:50) and anti-paxillin (ThermoScientific PA5-34910, 1:1000). Primary antibodies were visualised using Alexa Fluor-conjugated secondary antibodies (ThermoScientific, 1:500). To visualise the nucleus and the actin cytoskeleton, cells were also incubated with DAPI (ThermoScientific) and rhodamine phalloidin (ThermoScientific, 1:200), respectively. Monochrome images of each fluorophore were obtained at 40× magnification (NA, 1.3) using the EVOS FL2 Auto System (ThermoScientific). Image-based cell profiling was used to quantify the morphological profiles of individual cells. CellProfiler 2.4.0 (The Broad Institute) was used to align images, correct illumination and detect individual cells from images of the actin cytoskeleton. Measurements of shape or geometry and fluorescence integrated intensity from individual cells were obtained using built-in measurement modules of CellProfiler^58^ (v3.0). Focal adhesions were identified by paxillin staining and localisation to the cell periphery. The length of focal adhesions was measured based on the length of the major axis of an ellipse fitted to individual focal adhesions.

### Cell migration study

Nanopatterned glass coverslips were mounted in live-imaging chambers (Attofluor™, Thermo Fischer Scientific) and cells were seeded at 5000 cells/cm^2^ in a 1 ml suspension. Immediately after seeding the cells, the chambers were mounted on an EVOS FL Auto microscope equipped with an incubator stage. Brightfield images were captured at 5 min intervals for 24 h. Images were pre-processed with a wavelet filter to increase the contrast of the cell body. Finally, cell area and migration were tracked using the TrackMate^59^ plugin for FIJI.

### Transfection

Cells were first plated on conventional tissue culture polystyrene plates at 10000 cells/cm^2^. After overnight growth, the complete cell medium was replaced with medium without serum and antibiotics, to prepare cells for transfection. Cells were transfected with a total of 0.1 μg/cm^2^ of GFP-LifeAct and 0.4 μg/cm^2^ of either TfR-HaloTag^60^ (a gift from Matthew Kennedy at the University of Colorado, Denver), Paxillin-HaloTag^61^ or GFP-FAK (Addgene 50515). Plasmids were diluted in OptiMEM (ThermoFisher) and mixed with the FuGENE HD transfection reagent (Promega) at a ratio of 1:3.5 DNA (μg):transfection reagent. Cells were incubated with transfectants for 24 h before re-plating onto patterned coverslips for imaging. HaloTag labelling was performed by incubating cells for 20 min with Janelia Fluor 647^62^ (a generous gift from Luke Lavis, HHMI Janelia Farm) in complete growth medium at a final concentration of 200 nM.

### Dynamic FAK data

Cells transfected with GFP-FAK and tdTomato-LifeAct were grown for 24 h on nanopatterned PMMA. A laser scanning confocal microscope (Zeiss LSM 800) with on-stage incubation and focus lock was used to monitor cells over time. Time-lapse microscopy was performed using 63× magnification with oil immersion (NA, 1.4) at 5 min intervals for a total of 35 min. Cells were illuminated with a 488 nm laser to visualise GFP-FAK and a 561 nm laser to visualise tdTomato-LifeAct. The focal adhesion analysis server was used for the automated segmentation and analysis of dynamic focal adhesion properties^63,64^. The focal adhesion assembly and disassembly rates were calculated from stable focal adhesions (longevity > 20 min). Focal adhesions were measured from whole cells every 5 min for a total of 35 min.

### Stochastic optical reconstruction microscopy (STORM)

STORM was performed on a Zeiss Axio Observer microscope equipped with a 1.49 NA 100× objective. A 180 mW 638 nm laser (Vortran Stradus) was fibre coupled via a custom-built laser bed into the microscope using a modified Zeiss TIRF slider with an illumination footprint of ~75 μm, giving an estimated power density of 2 kW/cm^2^ after losses in the fibre and slider. A Definite Focus module was used to maintain focus during STORM image acquisition, which occurred over 5–20 min. Images were acquired using an Evolve Delta 512 EMCCD camera with a pixel size of 160 nm, an EM gain of 100 and an integration time of 18 ms for full frame and 10 ms for quarter frame crops. During STORM acquisition, reactivation of the fluorophores was controlled by modulating the power of a 408 nm laser (Vortran Stradus) on the same laser bed. Samples were mounted in an airtight silicone chamber in a STORM imaging buffer adapted from Olivier et al^65^. We imaged a single sample for 2–3 h before replacing the buffer; whenever buffer was changed, we allowed equilibration of the cell system for 30 min before imaging. Reference widefield images and confocal images were acquired of cells selected for STORM microscopy using the aforementioned camera or an LSM800 module attached to the same microscope body. STORM reconstructions were produced using the modified ThunderSTORM^66^ plugin for FIJI including the phasor fitting method with a pixel radius of 1. Detections below a photon threshold of 200 and below the sigma range were discarded.

### CLEM sample preparation

After STORM microscopy, samples were prepared for electron microscopy. The imaging buffer was replaced with water with minimal agitation of the sample, to preserve the fiducial beads that were in place. A secondary fixation in osmium tetroxide/glutaraldehyde was followed by dehydration in graded ethanol/water mixtures before final dehydration by critical point drying. Samples were sputter coated with 5 nm of gold–palladium and mounted for SEM imaging using a Hitachi SU8240 system in the SE mode and a 10 kV acceleration voltage. A conducting polymer solution was used to make contact between the sputtered substrate of the glass coverslip and a carrier piece of silicon, which minimised charging and image drifting during acquisition. STORM and SEM data were correlated using the *ec-CLEM* plugin for ICY^67^.

### Correlation between nanopatterns and STORM localisations

Prior to the acquisition of images using STORM, cells were imaged using brightfield illumination and 60× magnification. The brightfield images captured large fiducial markers (600 × 600 nm boxes) that denoted the position relative to the entire nanopatterned substrate. To ensure alignment to the fiducial markers within a 70 nm error from a true grid, brightfield images were transformed using an affine transformation. The resulting affine transformation matrix was then used to correct the STORM localisations, which were used in subsequent analyses. This methodology is detailed in Supplementary Figures 13–16 and Supplementary Note 3.

### Paxillin cluster analysis

Clusters of paxillin molecules were identified using the Hierarchical Density-based Spatial Clustering of Applications with Noise (HDBSCAN) algorithm and implementation^33^ for Python. The clustering algorithm allows the identification of clusters of varying sizes using a density-based approach. Essentially, clusters are defined by regions with a higher density that that of the surrounding space. To identify nascent clusters from paxillin molecules, we defined the following parameters in the HDBSCAN algorithm: minimum cluster size = 15 and minimum samples = 30 using the leaf cluster selection method. We produced images of the identified clusters from paxillin molecules (with and without nanopits, see Supplementary Figures 5 and 6) using the matplotlib^68^ package for Python. From these images, we segmented and extracted location and shape measurements from clusters and pits (such as diameter and area) using CellProfiler^58^ (v3.0). The identified clusters of paxillin were then used as nascent adhesions. The nearest neighbour distances between clusters and pits were measured using a custom code written in Python. We used the centroid coordinates of clusters and pits to measure d_cluster_, L_cluster–cluster_, L_cluster–pit_ and L_cluster–object_.

### Unsupervised machine learning

Nearest neighbour distances and paxillin cluster diameter were used as metrics for *k*-means clustering and principal component analysis. Data averaged from regions of interest were transformed by mean centring, followed by normalisation to the standard deviation before use. Both unsupervised machine learning algorithms were performed using the FactoMineR^69^ and factoextra^70^ packages for R. For k-means clustering, the optimal k number of clusters was obtained by maximisation of the average silhouette width (Supplementary Figure 17).

### Spatial statistics

We applied robust spatial statistics^35^ to determine the correlation and spacing between paxillin clusters and other clusters or nanopits. The spatial statistics analysis was carried out using the spatstat^71^ (v 1.63-3) package for R. Nascent adhesions identified from clusters of paxillin molecules were treated as point patterns, and nanopits were treated either as point patterns or spatial covariates. Multiple regions of interest (ROIs; sized 4 × 4 μm) from the same cell were analysed together as multiple views of the same object window. A total of n = 1607/1948/1958 paxillin clusters from 3 cells each on FLAT/SQ/NSQ were used in spatial statistics analyses. Intensity (number per unit area) was calculated from individual ROIs and is presented as an average value from multiple ROIs within one cell.

Analyses using summary statistics utilised border correction to restrict summary statistics to measurements that were completely within the ROI. The border correction excludes uncaptured relationships between clusters at the edge of the ROI and clusters outside of the ROIs. Empirical summary statistics were obtained by pooling together the summary statistics calculated on different cells from the same substrate. The spacing between paxillin clusters and nanopits was measured using the *J_cluster–pit_ (r)* function corrected for inhomogeneous intensity. The *J_cluster–pit_ (r)* function is defined as follows:

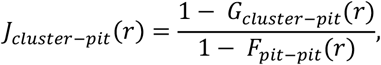

where *G_cluster-pit_ (r)* is the cumulative distribution function of the distance ≤ *r* between a typical paxillin cluster *x* at an arbitrary point *u*. The closest nanopit *x*^*pit*^ is defined as follows:

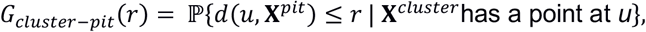

where *d*(*u*, **X**^*pit*^) = min {│|*u* − *x*^*pit*^|│: *x*^*pit*^ *ϵ* **X**^*pit*^}.

*F_pit–pit_ (r)* is the cumulative distribution function of the distance from an arbitrary location *u* to the nearest pit *x*^*pit*^:

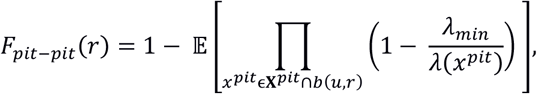

where *b*(*u*, *r*) defines the disc of radius *r* centred at an arbitrary location *u*.

Thus, the *J* function summarises the probability of obtaining objects close together compared with objects far apart. *J_cluster–pit_ (r)* > 1 denotes a regular spatial pattern, *J_cluster–pit_ (r)* < 1 denotes a clustered spatial pattern and *J_cluster–pit_ (r)* = 1 denotes independence between clusters and pits. An additional analysis of point patterns was performed (Supplementary Figures 8 and 9, Supplementary Tables 1 and 2) and is described in detail in Supplementary Note 2.

### Single-particle tracking (SPT) microscopy

Cells transfected with HaloTag fusions were labelled with photoactivatable Janelia Fluor dyes in warm media for 20 min prior to washing and mounting in live-imaging buffer (Thermo). For imaging, we used the setup described above for STORM. Laser and camera triggering were controlled by an Arduino microcontroller, to activate a subset of molecules with the 408 nm laser and capture their movement for 30 s. We used a stroboscopic illumination scheme to limit blurring of the emitting probes, with 5 ms illumination at the start of a 10 ms camera integration time. Image acquisition was performed using the setup described for STORM. Data were processed using the TrackMate^59^ plugin for FIJI. Three ROIs of 8 × 8 μm were selected from both paxillin and TfR STORM data sets. In each case, the acquisition time per frame was 18–20 ms. The number of frames was halved by adding together pairs of images to improve the signal-to-noise ratio, yielding a final frame count for each ROI of up to 7500 frames, with an effective acquisition time of 0.036 s per frame. Mobile particles or spots were detected in TrackMate using a blob diameter of 0.5 μm, with a corresponding linking distance and gap-closing maximum distance of 0.5 μm. A gap-closing maximum frame gap of 1 frame was used. Spot size filters were also used to eliminate artefacts (spots smaller than 0.1 μm were rejected). Track filters were used to reject background (minimum number of spots per track, 9; median velocity, above 1.0 and below 7.0).

Track metrics (duration, confinement ratio, velocity and rate of linear progression) were obtained using TrackMate. For further analysis, the data were filtered to exclude tracks with zero displacement (immobile tracks). The diffusion coefficient and alpha coefficient were estimated from filtered data using the msdanalyzer^72^ toolbox for MATLAB. From the first 50% of the mean of time-averaged ensemble MSD, we obtained a linear fit to estimate the diffusion coefficient under different substrates. The diffusion coefficient was then used to calculate the value of α, which indicates the dependency on the MSD 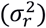 and time (*t*):

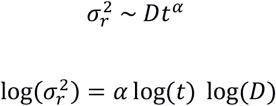

The estimation of the α values was carried out by fitting a straight line against 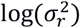 vs. log(*t*). Only tracks with a goodness-of-fit measure *R*^2^ ≥ 80% were included in the estimation of α. Thereafter, we determined the number of diffusive substates, corresponding diffusion coefficients and occupancy of tracks under each substrate condition using a Bayesian variational approach implemented in the vbSPT^39^ toolbox for MATLAB.

### Correlating nanopattern and SPT data

For single-particle tracking, additional accuracy in the alignment of microscopy images to the true nanopattern was achieved by first drying fluorescent polystyrene beads (0.1 μm diameter, diluted to 50000:1 in water; ThermoFisher) on the nanopatterned substrate. As the evaporating solution moved across the pattern, a sparse number of beads were pushed into random points of the nanopattern, which provided fiducial markers built into the nanopattern for correlating nanopattern location with microscopy data. Robust matching between the polystyrene beads and the nanopits was performed using RANSAC optimisation. Beads that were found to be inliers were then used as patterns to match the underlying nanopatterns, yielding a second affine transformation. In summary, SPT data were aligned to the nanopatterns through two affine transformation matrices. This methodology is detailed in Supplementary Figures 13–16 and Supplementary Note 3.

### Statistics, visualisation and software

Statistical tests were performed using the statistical software R^73^ (v3.5.1) through its graphical interface RStudio (v1.2.1114). Excluding the statistical tests used for spatial statistics, a one-way or two-way ANOVA (specified in the figure legend) with Tukey’s post hoc test for multiple comparisons were used. Scatterplots, boxplots, histograms, density plots and line plots were generated using the ggplot2 (v3.3.0.9000) package for R. Visualisation of diffusion tracks with bounding boxes was performed using the matplotlib^68^ package for Python^74^ (v. 3.6).

## Acknowledgements

This study was funded by the European Research Council FAKIR 648892 Consolidator Award. MFAC thanks financial support from the University of Glasgow MG Dunlop Bequest, College of Science and Engineering Scholarship, and the FAKIR consolidator award. Thanks to the microscopy community on Twitter for detailed advice on super resolution microscopy, particularly Dr Christophe Leterrier (@christlet) and Dr Riccardo Henriques (@riccardohenriques). MFAC and PMR thanks Rachel Duckhouse (@rachelduckhouse), our artist in residence (2018), for stimulating conversations on disorder, symmetry, and functionality.

## Author Contributions

MFAC: Investigation, data curation, visualisation, formal analysis, writing (original draft preparation, review and editing)

CS: Data curation, visualisation, formal analysis, writing (original draft preparation, review and editing)

PMR: Conceptualisation, methodology, investigation, data curation, visualisation, formal analysis, writing (original draft preparation, review and editing)

NG: Conceptualisation, methodology, writing (review and editing), funding acquisition, supervision

## References

1. Curtis, A. S. G. et al. Cells react to nanoscale order and symmetry in their surroundings. IEEE Trans. Nanobioscience 3, 61–65 (2004).

2. Dalby, M. J. et al. The control of human mesenchymal cell differentiation using nanoscale symmetry and disorder. Nat. Mater. 6, 997–1003 (2007).

3. McMurray, R. J. et al. Nanoscale surfaces for the long-term maintenance of mesenchymal stem cell phenotype and multipotency. Nat. Mater. 10, 637–644 (2011).

4. Tsimbouri, P. et al. Nanotopographical effects on mesenchymal stem cell morphology and phenotype. J. Cell. Biochem. 115, 380–390 (2014).

5. Tsimbouri, P. M. et al. Using nanotopography and metabolomics to identify biochemical effectors of multipotency. ACS Nano 6, 10239–10249 (2012).

6. Yang, J. et al. Nanotopographical induction of osteogenesis through adhesion, bone morphogenic protein cosignaling, and regulation of MicroRNAs. ACS Nano 8, 9941–9953 (2014).

7. Ngandu Mpoyi, E. et al. Protein adsorption as a key mediator in the nanotopographical control of cell behavior. ACS Nano 10, 6638–6647 (2016).

8. Biggs, M. J. P. et al. Interactions with nanoscale topography: adhesion quantification and signal transduction in cells of osteogenic and multipotent lineage. J. Biomed. Mater. Res. A 91, 195–208 (2009).

9. Lee, L. C. Y. et al. Nanotopography controls cell cycle changes involved with skeletal stem cell self-renewal and multipotency. Biomaterials 116, 10–20 (2017).

10. Cutiongco, M. F. A., Jensen, B. S., Reynolds, P. M. & Gadegaard, N. Predicting gene expression using morphological cell responses to nanotopography. Nat. Commun. 11, 1384 (2020).

11. Allan, C. et al. Osteoblast response to disordered nanotopography. J. Tissue Eng. 9, 204173141878409–7 (2018).

12. Kantawong, F. et al. Whole proteome analysis of osteoprogenitor differentiation induced by disordered nanotopography and mediated by ERK signalling. Biomaterials 30, 4723–4731 (2009).

13. Dalby, M. J. et al. Nanomechanotransduction and interphase nuclear organization influence on genomic control. J. Cell. Biochem. 102, 1234–1244 (2007).

14. Biggs, M. J. P. et al. The use of nanoscale topography to modulate the dynamics of adhesion formation in primary osteoblasts and ERK/MAPK signalling in STRO-1+ enriched skeletal stem cells. Biomaterials 30, 5094–5103 (2009).

15. Ross, E. A. et al. Nanotopography reveals metabolites that maintain the immunosuppressive phenotype of mesenchymal stem cells. bioRxiv 603332 (2019) doi:10.1101/603332.

16. Dalby, M. J., Gadegaard, N. & Oreffo, R. O. C. Harnessing nanotopography and integrin–matrix interactions to influence stem cell fate. Nat. Publ. Group 13, 558–569 (2014).

17. Dalby, M. J. et al. Nanotopographical stimulation of mechanotransduction and changes in interphase centromere positioning. J. Cell. Biochem. 100, 326–338 (2007).

18. Gallagher, J. O., McGhee, K. F., Wilkinson, C. D. W. & Riehle, M. O. Interaction of animal cells with ordered nanotopography. IEEE Trans. Nanobioscience 99, 24–28 (2002).

19. Dalby, M. J., Gadegaard, N., Riehle, M. O., Wilkinson, C. D. W. & Curtis, A. S. G. Investigating filopodia sensing using arrays of defined nano-pits down to 35 nm diameter in size. Int. J. Biochem. Cell Biol. 36, 2005–2015 (2004).

20. Lamers, E. et al. The influence of nanoscale grooved substrates on osteoblast behavior and extracellular matrix deposition. Biomaterials 31, 3307–3316 (2010).

21. Park, J., Bauer, S., von der Mark, K. & Schmuki, P. Nanosize and vitality: TiO 2Nanotube diameter directs cell fate. Nano Lett. 7, 1686–1691 (2007).

22. Lim, J. et al. Constrained adherable area of nanotopographic surfaces promotes cell migration through the regulation of focal adhesion via focal adhesion Kinase/Rac1 activation. ACS Appl. Mater. Interfaces 10, 14331–14341 (2018).

23. Cavalcanti-Adam, E. A. et al. Cell spreading and focal adhesion dynamics are regulated by spacing of integrin ligands. Biophys. J. 92, 2964–2974 (2007).

24. Cavalcanti-Adam, E. A., Aydin, D., Hirschfeld-Warneken, V. C. & Spatz, J. P. Cell adhesion and response to synthetic nanopatterned environments by steering receptor clustering and spatial location. HFSP J. 2, 276–285 (2008).

25. Huang, J. et al. Impact of order and disorder in RGD nanopatterns on cell adhesion. Nano Lett. 9, 1111–1116 (2009).

26. Xie, Y. et al. Quantitative profiling of spreading-coupled protein tyrosine phosphorylation in migratory cells. Sci. Rep. 1–10 (2016) doi:10.1038/srep31811.

27. Yang, B. et al. Mechanosensing controlled directly by tyrosine kinases. Nano Lett. 16, 5951–5961 (2016).

28. Jiang, G., Huang, A. H., Cai, Y., Tanase, M. & Sheetz, M. P. Rigidity sensing at the leading edge through alphavbeta3 integrins and RPTPalpha. Biophysj 90, 1804–1809 (2006).

29. Ghassemi, S. et al. Cells test substrate rigidity by local contractions on submicrometer pillars. Proc. Natl. Acad. Sci. 109, 5328–5333 (2012).

30. Turner, C. E. Paxillin and focal adhesion signalling. Nat. Cell Biol. 2, E231–E236 (2000).

31. Deakin, N. O. & Turner, C. E. Paxillin comes of age. J. Cell Sci. 121, 2435–2444 (2008).

32. Changede, R., Xu, X., Margadant, F. & Sheetz, M. P. Nascent integrin adhesions form on all matrix rigidities after integrin activation. Dev. Cell 35, 614–621 (2015).

33. Campello, R. J. G. B., Moulavi, D. & Sander, J. Density-based clustering based on hierarchical density estimates. in Advances in knowledge discovery and data mining (eds. Pei, J., Tseng, V. S., Cao, L., Motoda, H. & Xu, G.) 160–172 (Springer Berlin Heidelberg, 2013).

34. Changede, R., Cai, H., Wind, S. J. & Sheetz, M. P. Integrin nanoclusters can bridge thin matrix fibres to form cell–matrix adhesions. Nat. Mater. 18, 1366–1375 (2019).

35. Baddeley, A., Rubak, E. & Turner, R. Spatial point patterns: methodology and applications with R. (CRC Press, Taylor & Francis Group, 2016).

36. Shibata, A. C. E. et al. Archipelago architecture of the focal adhesion: membrane molecules freely enter and exit from the focal adhesion zone. Cytoskelet. Hoboken NJ 69, 380–392 (2012).

37. Hu, Y.-L. et al. FAK and paxillin dynamics at focal adhesions in the protrusions of migrating cells. Sci. Rep. 4, (2014).

38. Digman, M. A., Brown, C. M., Horwitz, A. R., Mantulin, W. W. & Gratton, E. Paxillin dynamics measured during adhesion assembly and disassembly by correlation spectroscopy. Biophys. J. 94, 2819–2831 (2008).

39. Persson, F., Lindén, M., Unoson, C. & Elf, J. Extracting intracellular diffusive states and transition rates from single-molecule tracking data. Nat. Methods 10, 265–269 (2013).

40. Kusumi, A. et al. Dynamic Organizing Principles of the Plasma Membrane that Regulate Signal Transduction: Commemorating the Fortieth Anniversary of Singer and Nicolson’s Fluid-Mosaic Model. Annu. Rev. Cell Dev. Biol. 28, 215–250 (2012).

41. Fujiwara, T., Ritchie, K., Murakoshi, H., Jacobson, K. & Kusumi, A. Phospholipids undergo hop diffusion in compartmentalized cell membrane. J. Cell Biol. 157, 1071–1081 (2002).

42. Fujiwara, T. K. et al. Confined diffusion of transmembrane proteins and lipids induced by the same actin meshwork lining the plasma membrane. Mol. Biol. Cell 27, 1101–1119 (2016).

43. Murase, K. et al. Ultrafine Membrane Compartments for Molecular Diffusion as Revealed by Single Molecule Techniques. Biophys. J. 86, 4075–4093 (2004).

44. Saxton, M. J. Anomalous diffusion due to obstacles: a Monte Carlo study. Biophys. J. 66, 394–401 (1994).

45. Kalay, Z., Fujiwara, T. K., Otaka, A. & Kusumi, A. Lateral diffusion in a discrete fluid membrane with immobile particles. Phys. Rev. E Stat. Nonlin. Soft Matter Phys. 89, 022724 (2014).

46. Werner, J. H. et al. Formation and Dynamics of Supported Phospholipid Membranes on a Periodic Nanotextured Substrate. Langmuir 25, 2986–2993 (2009).

47. Curtis, A. S. G., Dalby, M. J. & Gadegaard, N. Nanoprinting onto cells. J. R. Soc. Interface 3, 393–398 (2006).

48. Sansen, T. et al. Mapping Cell Membrane Organization and Dynamics Using Soft Nano-Imprint Lithography. ACS Appl. Mater. Interfaces (2020) doi:10.1021/acsami.0c05432.

49. Minton, A. P. Confinement as a determinant of macromolecular structure and reactivity. Biophys. J. 63, 1090–1100 (1992).

50. Zhou, H.-X., Rivas, G. & Minton, A. P. Macromolecular crowding and confinement: biochemical, biophysical, and potential physiological consequences. Annu. Rev. Biophys. 37, 375–397 (2008).

51. Guigas, G. & Weiss, M. Sampling the Cell with Anomalous Diffusion—The Discovery of Slowness. Biophys. J. 94, 90–94 (2008).

52. Cheng, B. et al. Nanoscale integrin cluster dynamics controls cellular mechanosensing via FAKY397 phosphorylation. Sci. Adv. 6, eaax1909 (2020).

53. Salasznyk, R. M., Klees, R. F., Williams, W. A., Boskey, A. & Plopper, G. E. Focal adhesion kinase signaling pathways regulate the osteogenic differentiation of human mesenchymal stem cells. Exp. Cell Res. 313, 22–37 (2007).

54. Castillo, A. B. et al. Focal Adhesion Kinase Plays a Role in Osteoblast Mechanotransduction In Vitro but Does Not Affect Load-Induced Bone Formation In Vivo. PLOS ONE 7, e43291 (2012).

55. Grashoff, C. et al. Measuring mechanical tension across vinculin reveals regulation of focal adhesion dynamics. Nature 466, 263–266 (2010).

56. Geiger, B., Spatz, J. P. & Bershadsky, A. D. Environmental sensing through focal adhesions. Nat. Rev. Mol. Cell Biol. 10, 21–33 (2009).

57. Huethorst, E. et al. Customizable, engineered substrates for rapid screening of cellular cues. Biofabrication 12, 025009 (2020).

58. McQuin, C. et al. CellProfiler 3.0: Next-generation image processing for biology. PLoS Biol. 16, e2005970–17 (2018).

59. Tinevez, J.-Y. et al. TrackMate: An open and extensible platform for single-particle tracking. Methods 115, 80–90 (2017).

60. Bowen, A. B., Bourke, A. M., Hiester, B. G., Hanus, C. & Kennedy, M. J. Golgi-independent secretory trafficking through recycling endosomes in neuronal dendrites and spines. eLife 6, e27362 (2017).

61. Lee, K. et al. Functional Hierarchy of Redundant Actin Assembly Factors Revealed by Fine-Grained Registration of Intrinsic Image Fluctuations. Cell Syst. 1, 37–50 (2015).

62. Grimm, J. B. et al. A general method to improve fluorophores for live-cell and single-molecule microscopy. Nat. Methods 12, 244–250 (2015).

63. Berginski, M. E. & Gomez, S. M. The Focal Adhesion Analysis Server: a web tool for analyzing focal adhesion dynamics. F1000Research 2, 68 (2013).

64. Berginski, M. E., Vitriol, E. A., Hahn, K. M. & Gomez, S. M. High-resolution quantification of focal adhesion spatiotemporal dynamics in living cells. PLoS ONE 6, e22025–13 (2011).

65. Olivier, N., Keller, D., Gönczy, P. & Manley, S. Resolution doubling in 3D-STORM imaging through improved buffers. PLOS ONE 8, 1–9 (2013).

66. Ovesný, M., Křížek, P., Borkovec, J., Švindrych, Z. & Hagen, G. M. ThunderSTORM: a comprehensive ImageJ plug-in for PALM and STORM data analysis and super-resolution imaging. Bioinformatics 30, 2389–2390 (2014).

67. Paul-Gilloteaux, P. et al. eC-CLEM: flexible multidimensional registration software for correlative microscopies. Nat. Methods 14, 102–103 (2017).

68. J. D. Hunter. Matplotlib: A 2D Graphics Environment. Comput. Sci. Eng. 9, 90–95 (2007).

69. Lê, S., Josse, J. & Husson, F. FactoMineR: An R Package for Multivariate Analysis. J. Stat. Softw. Vol 1 Issue 1 2008 (2008).

70. Kassambara, A. & Mundt, F. factoextra: Extract and visualize the results of multivariate data analyses. https://CRAN.R-project.org/package=factoextra (2017).

71. Baddeley, A. & Turner, R. spatstat: An R Package for Analyzing Spatial Point Patterns. J. Stat. Softw. 12, (2005).

72. Tarantino, N. et al. TNF and IL-1 exhibit distinct ubiquitin requirements for inducing NEMO–IKK supramolecular structures. J. Cell Biol. 204, 231–245 (2014).

73. Team, R. C. R: A language and environment for statistical computing. (2018).

74. Oliphant, T. E. Python for scientific computing. Comput. Sci. Eng. 9, 10–20.

