## Supplementary Information for "Nanotopography controls single-molecule mobility to determine overall cell fate"

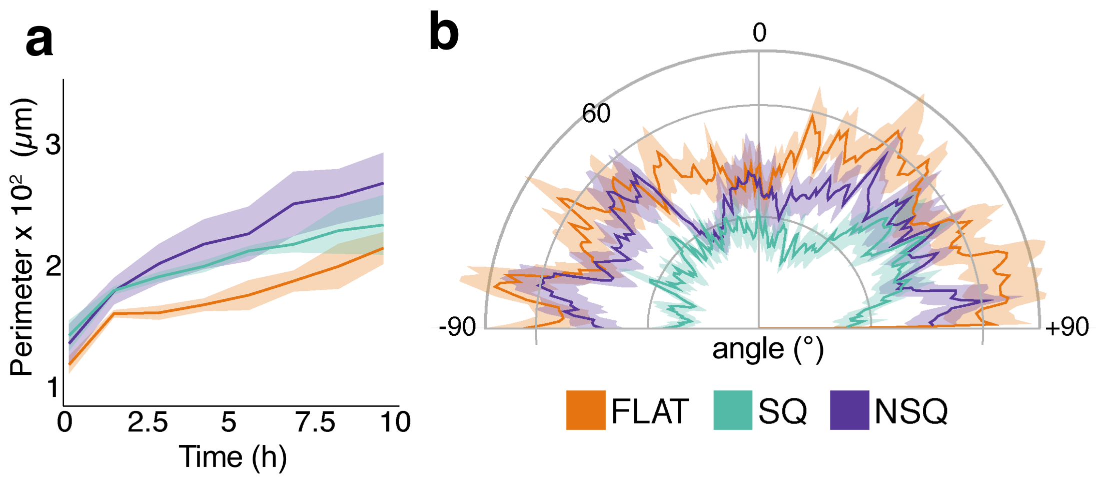


**Supplementary Figure 1. Nanopatterns change form factor and focal adhesion of cells.** a) Form factor of cells on nanopatterns measured over time. Form factor was calculated as the ratio of the area to the perimeter of each cell, with form factor = 1 indicating a perfect circle. Data were obtained from three independent experiments for a total of n = 16/11/11 for FLAT/SQ/NSQ. b) Orientation of focal adhesions. OrientationJ[1] for Fiji was used to determine the orientation of individual focal adhesions from entire cells. Data were obtained from one independent experiment for a total of n = 45/42/40 for FLAT/SQ/NSQ. The lines and the corresponding shaded areas indicate the mean ± standard deviation.

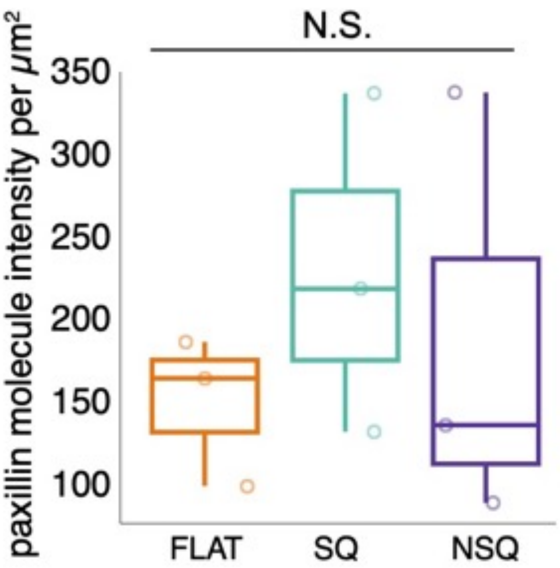


**Supplementary Figure 2.** Intensity (count per $\mu m^2$ or $\lambda$) of paxillin molecules. NS denotes the absence of statistical significance for all pairwise comparisons. Statistical analysis was performed using one-way ANOVA with Tukey's post-hoc test. The points indicate intensity measurements obtained from individual cells.

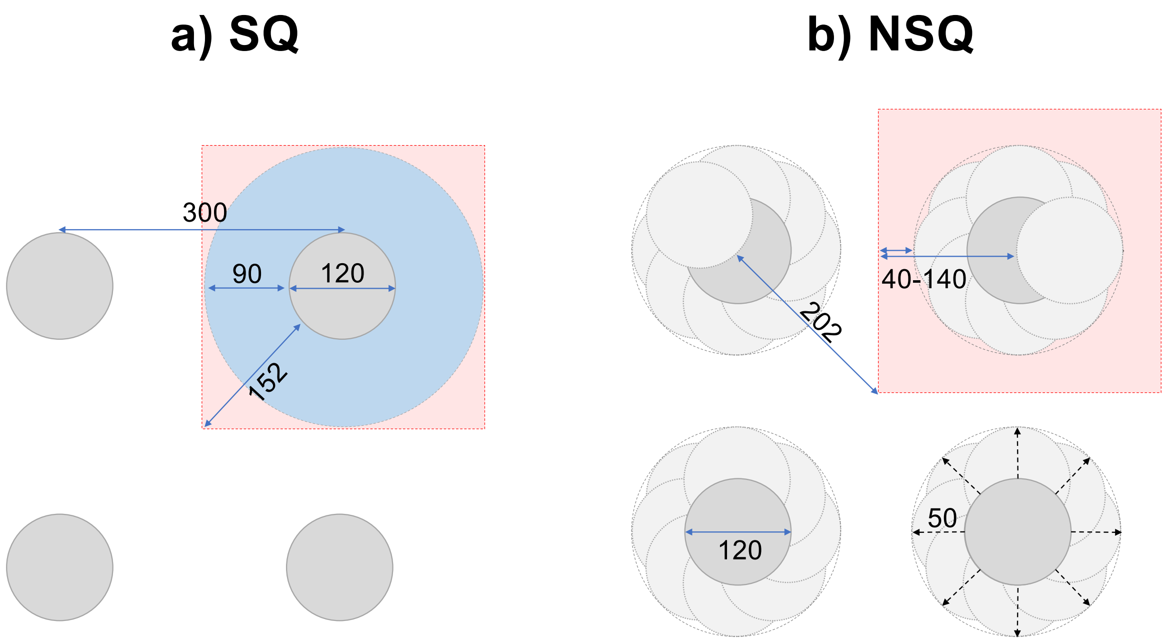


**Supplementary Figure 3.** Schematic representation of SQ and NSQ nanopit geometries. All pits are 120 nm in diameter and adjacent pits are spaced at a 300 nm pitch (centre to centre). a) On SQ, the halfway distances between neighbouring nanopits ranged from 90 nm (for adjacent nanopits, or first-nearest neighbours) to a maximum of 152 nm (for diagonally opposite nanopits, or second-nearest neighbours). b) On NSQ, nanopits were displaced from an SQ arrangement by +/– 50 nm in any direction, resulting in a variation in the halfway distance between nearest nanopits. The halfway distance to the nearest nanopit edge lay between 40–140 nm (between first-nearest neighbours) and 202 nm (between second-nearest neighbours).

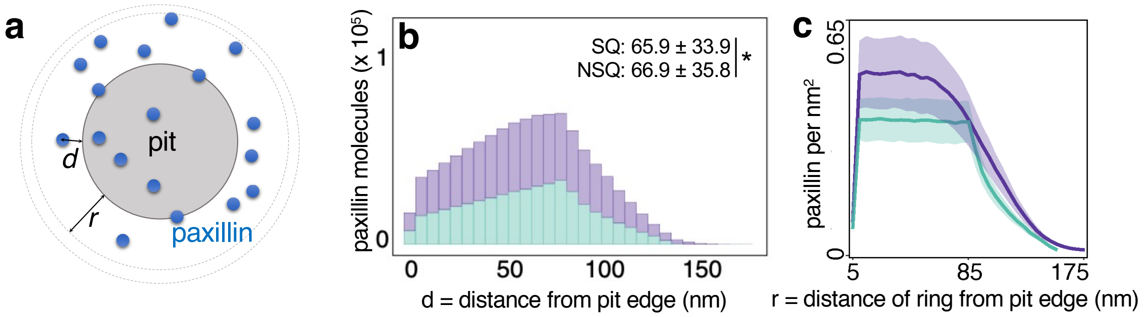


**Supplementary Figure 4.** Nanopits do not nucleate the aggregation of paxillin molecules. a) Schematic representation of the analysis of the paxillin–nanopit interaction. The distance from each paxillin molecule to the edge of the nearest nanopit (denoted as *d*) was measured, and the number of paxillin in each ring of 10 nm thickness was assessed with increasing distance to the nanopit edge (denoted as *r*). b) Distance from the paxillin molecule to the edge of the nearest nanopit (*d*). Data are binned every 5 nm. c) Paxillin density as a function of the distance from the pit edge (*r*). Line plots show the mean ± confidence interval, boxplots show the median and interquartile range with minima and maxima at the whiskers and the dots indicate individual data points. n = 18/18 regions of interest from three cells each on SQ/NSQ. * Denotes statistical significance between indicated pairs, as assessed using one-way ANOVA with Tukey's post-hoc test.

**Supplementary Note 1**

Testing the nucleating effect of nanopits

We analysed data from super-resolution microscopy to test the hypothesis that nanopit edges nucleate focal adhesion formation. To do this, we considered the relationship between the distance from paxillin to nanopit edges and this phenomenon. If nanopit edges nucleated paxillin molecule aggregation (Supplementary Figure 4a), then paxillin molecules should show a higher likelihood to be found nearer to a nanopit edge. Under this assumption, the shortest distance between a paxillin molecule and the nearest nanopit edge would be significantly shorter than the halfway distances between nearest neighbouring pits; i.e., paxillin molecules that lie halfway between first- or second-nearest neighbouring pits show no preference for location near either of the nanopits.

This phenomenon would be easily apparent on SQ, where the halfway distances between nearest neighbouring nanopits only varied between 90 and 152 nm (Supplementary Figure 3). However, the distribution of distances between a paxillin molecule and the nearest neighbouring nanopit edge was almost flat between 10 and 100 nm (Supplementary Figure 4b). In fact, paxillin molecules were found on average to be as far away as 65.9 nm from the nearest nanopit edge. This indicated that paxillin molecules are less likely to be found near nanopit edges than between nanopits. Although the distances between nanopits on NSQ were highly variable and depended on the local disorder in the nanopits (Supplementary Figure 3), we observed a similar trend. We found no clear peak in the distribution of the distance between a paxillin molecule and the nearest neighbouring nanopit edge. Instead, we found an even flatter distribution of distances between paxillin and the nearest neighbouring nanopit edge compared with SQ.

We further analysed the dataset in terms of density. We calculated paxillin density in 10 nm rings with increasing distance from the nearest nanopit edge (Supplementary Figure 4a). Under the assumption that nanopits nucleate focal adhesion formation, we would expect paxillin density to slope downward, with the maximum found near the nanopit edge. Here again our results contradicted this hypothesis (Supplementary Figure 4c). Instead of a downward slope, we observed no change in paxillin density with respect to distance from the nanopit edge (regardless of the nanopit arrangement). Our results demonstrate that paxillin density does not depend on the distance to the nanopit edge.

Hence, our data contradicts the hypothesis that nanopits nucleate the aggregation of focal-adhesion components, such as paxillin.

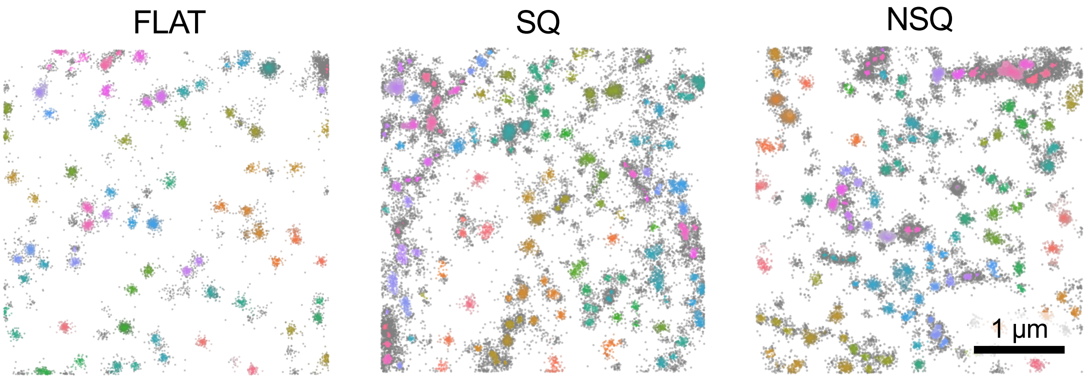


**Supplementary Figure 5.** Paxillin clustering on different nanopatterns. Representative images of clusters identified from individual paxillin molecules. The colour scheme denotes the membership of individual paxillin molecules to the same cluster, with small grey dots indicating paxillin molecules that were not clustered. Clustering was performed using the HDBSCAN algorithm[2] (see Materials and Methods).

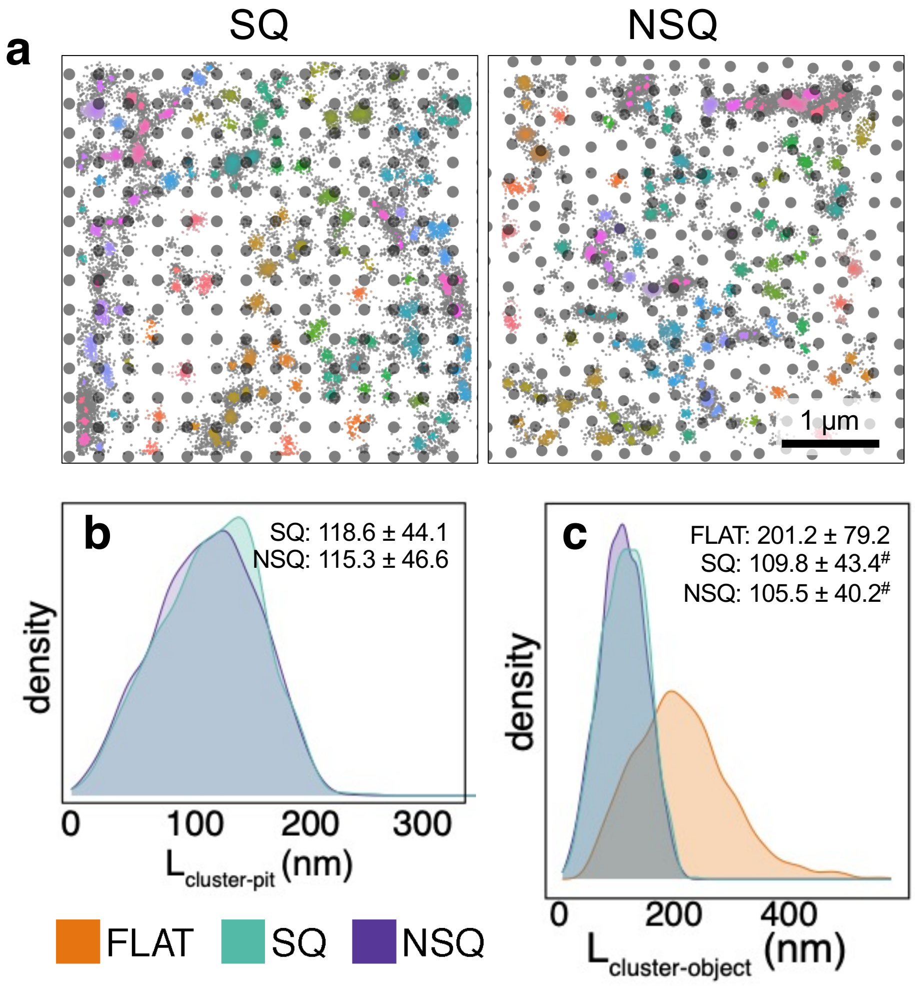


**Supplementary Figure 6.** Interaction between paxillin clusters and nanopits. a) Representative images of the clusters identified from paxillin molecules overlaid with the nanopit array. Large grey dots denote nanopits. The colour scheme indicates the membership of individual paxillin molecules to the same cluster, with small grey dots denoting paxillin molecules that were not clustered. b) Distribution of the distance between a cluster and the nearest neighbouring pit ($L_{cluster–pit}$) across different substrates. c) Distribution of the distance between a cluster and the nearest neighbouring object (either a cluster or a pit, denoted as $L_{cluster–object}$) across different substrates. For FLAT, $L_{cluster–pit}$ = $L_{cluster–cluster}$. n = 15/13 regions analysed for SQ/NSQ obtained from three independent experiments. # Denotes statistical significance against FLAT, as assessed using one-way ANOVA with Tukey's post hoc test.

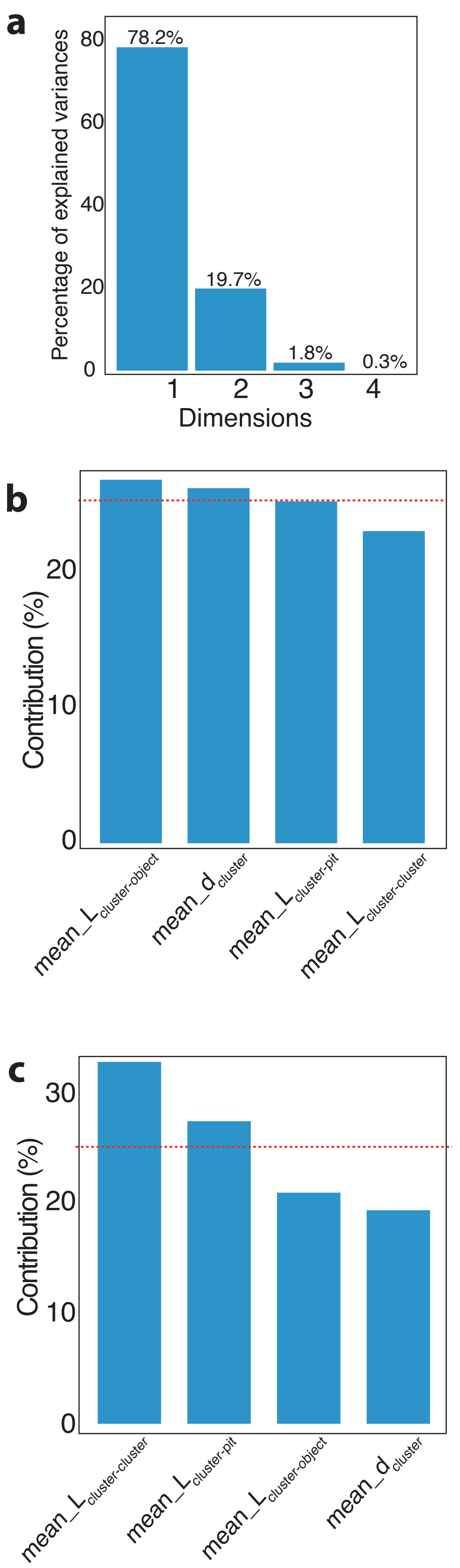


**Supplementary Figure 7.** Principal component analysis. Principal component analysis was performed using mean and standard deviation (SD) measurements of paxillin cluster diameter ($d_{cluster}$) and nearest neighbour distances ($L_{cluster–cluster}$, $L_{cluster–pit}$ $L_{cluster–object}$) measured from FLAT, SQ and NSQ. a) The variance explained by each dimension of the principal components. b–c) The contribution of each measurement to the variance explained by the first two dimensions of the principal components.

**Supplementary Table 1.** Spatial pattern of paxillin clusters across nanopatterns. The Berman test is a non-parametric hypothesis test that is used to determine if the intensity of the point process shows complete spatial randomness or no relationship between clusters (null hypothesis). The Berman test is based on the cumulative distribution function of the intensity. Further hypothesis testing of the correlation and spacing of paxillin clusters was performed using the maximum absolute deviation (MAD) test. The MAD test is used to determine if the point process shows complete spatial randomness (null hypothesis).

| **Topography** | **Cell number** | **Berman test for intensity $\lambda_{cluster}$** | **MAD test for $L_{cluster–cluster}(r)$** | **MAD test for $G_{cluster–cluster}(r)$** | **MAD test for $J_{cluster–cluster}(r)$** |
|---|---|---|---|---|---|
| NSQ | 1 | Z2 = 34.689, *P*-value < 2.2e–16 | mad = 0.054076, rank = 1, *P*-value = 0.01 | mad = 0.0072573, rank = 100, *P*-value = 1 | mad = 0.0082435, rank = 100, *P*-value = 1 |
| NSQ | 2 | Z2 = 35.021, *P*-value < 2.2e–16 | mad = 0.047693, rank = 1, *P*-value = 0.01 | mad = 0.0023668, rank = 100, *P*-value = 1 | mad = 0.0026198, rank = 100, *P*-value = 1 |
| NSQ | 3 | Z2 = 26.058, *P*-value < 2.2e–16 | mad = 0.049896, rank = 1, *P*-value = 0.01 | mad = 2.6714e-07, rank = 100, *P*-value = 1 | mad = 3.8477e-07, rank = 100, *P*-value = 1 |
| SQ | 1 | Z2 = 32.804, *P*-value < 2.2e–16 | mad = 0.030599, rank = 1, *P*-value = 0.01 | mad = 0.089368, rank = 100, *P*-value = 1 | mad = 0.1054, rank = 87, *P*-value = 0.87 |
| SQ | 2 | Z2 = 26.961, *P*-value < 2.2e–16 | mad = 0.058971, rank = 1, *P*-value = 0.01 | mad = 1.2807e-07, rank = 100, *P*-value = 1 | mad = 1.3696e-07, rank = 100, *P*-value = 1 |
| SQ | 3 | Z2 = 32.704, *P*-value < 2.2e–16 | mad = 0.027399, rank = 100, *P*-value = 1 | mad = 1.1166e-05, rank = 100, *P*-value = 1 | mad = 1.4616e-05, rank = 100, *P*-value = 1 |
| FLAT | 1 | Z2 = 35.597, *P*-value < 2.2e–16 | mad = 0.038699, rank = 1, *P*-value = 0.01 | mad = 3.2182e-06, rank = 100, *P*-value = 1 | mad = 4.5244e-06, rank = 100, *P*-value = 1 |
| FLAT | 2 | Z2 = 35.151, *P*-value < 2.2e–16 | mad = 0.057216, rank = 1, *P*-value = 0.01 | mad = 0.032266, rank = 100, *P*-value = 1 | mad = 0.038317, rank = 44, *P*-value = 0.44 |
| FLAT | 3 | Z2 = 27.295, *P*-value < 2.2e–16 | mad = 0.062121, rank = 1, *P*-value = 0.01 | mad = 0.07109, rank = 100, *P*-value = 1 | mad = 0.088724, rank = 99, *P*-value = 0.99 |

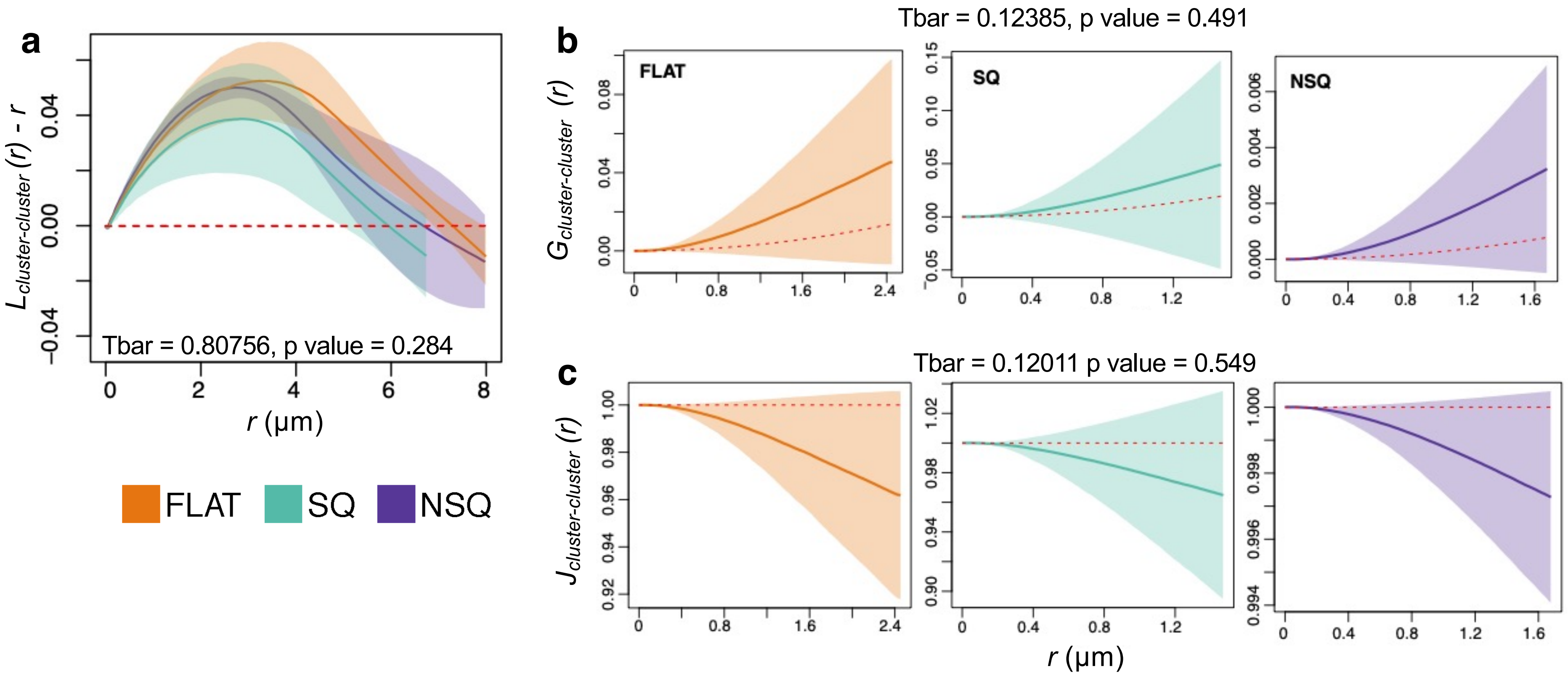


**Supplementary Figure 8.** Spatial distribution of paxillin clusters across nanopatterns. We calculated summary statistics that describe the correlation and spacing between paxillin clusters. a) The $L_{cluster\text{–}cluster}$ *(r)* function describes the spatial correlation between paxillin clusters. $L_{cluster\text{–}cluster}$ *(r)* > 0 indicates spatial clustering, while $L_{cluster\text{–}cluster}$ *(r)* < 0 indicates the spatial regularity of paxillin clusters. b) The G function describes the nearest neighbour distances between paxillin clusters. $G_{cluster\text{–}cluster}$ *(r)* values above the line representing a random process denote aggregation of paxillin clusters at the spatial scale *r*. Conversely, $G_{cluster\text{–}cluster}$ *(r)* values below the line representing a random process indicate a regular pattern at the spatial scale *r.* c) The J function describes the probability of finding paxillin clusters close together vs. those far apart. $J_{cluster\text{–}cluster}$ *(r)* > 1 denotes regularity and $J_{cluster\text{–}cluster}$ *(r)* < 1 denotes aggregation. The lines indicate the empirical summary function estimated from the dataset, while shaded areas represent the confidence interval. The red line denotes expected values for a random point process. We present the test statistic Tbar and *P* value from the permutation test of the summary statistic.

**Supplementary Table 2.** Dependence of the spatial pattern of paxillin cluster on nanopatterns. The Berman test was used to determine if the intensity of the point process depends on the location of the nanopits (alternative hypothesis). Further hypothesis testing on the correlation and spacing of paxillin clusters was performed using the maximum absolute deviation (MAD) test. The MAD test for multiple objects is used to determine if the point process shows spatial randomness of, and independence between, the location of paxillin clusters and nanopits (null hypothesis).

| Topography | Cell number | Berman test for $\lambda_{cluster-pit}$ | MAD test for $L_{cluster-pit}(r)$ | MAD test for $G_{cluster-pit}(r)$ | MAD test for $J_{cluster-pit}(r)$ |
|---|---|---|---|---|---|
| NSQ | 1 | Z2 = –1.9355, *P*-value = 0.05293 | mad = 0.031536, rank = 1, *P*-value = 0.01 | mad = 0.16467, rank = 1, *P*-value = 0.01 | mad = 0.28307, rank = 1, *P*-value = 0.01 |
| NSQ | 2 | Z2 = –1.6716, *P*-value = 0.09461 | mad = 0.021885, rank = 100, *P*-value = 1 | mad = 0.19512, rank = 1, *P*-value = 0.01 | mad = 0.25513, rank = 1, *P*-value = 0.01 |
| NSQ | 3 | Z2 = –0.65664, *P*-value = 0.5114 | mad = 0.059125, rank = 1, *P*-value = 0.01 | mad = 0.17771, rank = 1, *P*-value = 0.01 | mad = 0.2794, rank = 1, *P*-value = 0.01 |
| SQ | 1 | Z2 = –1.7277, *P*-value = 0.08405 | mad = 0.012724, rank = 100, *P*-value = 1 | mad = 0.24161, rank = 1, *P*-value = 0.01 | mad = 0.28278, rank = 1, *P*-value = 0.01 |
| SQ | 2 | Z2 = 1.6382, *P*-value = 0.1014 | mad = 0.027415, rank = 100, *P*-value = 1 | mad = 0.17777, rank = 1, *P*-value = 0.01 | mad = 0.26685, rank = 1, *P*-value = 0.01 |
| SQ | 3 | Z2 = 7.0362, *P*-value = 1.975e–12 | mad = 0.039009, rank = 100, *P*-value = 1 | mad = 0.27852, rank = 1, *P*-value = 0.01 | mad = 0.30387, rank = 1, *P*-value = 0.01 |

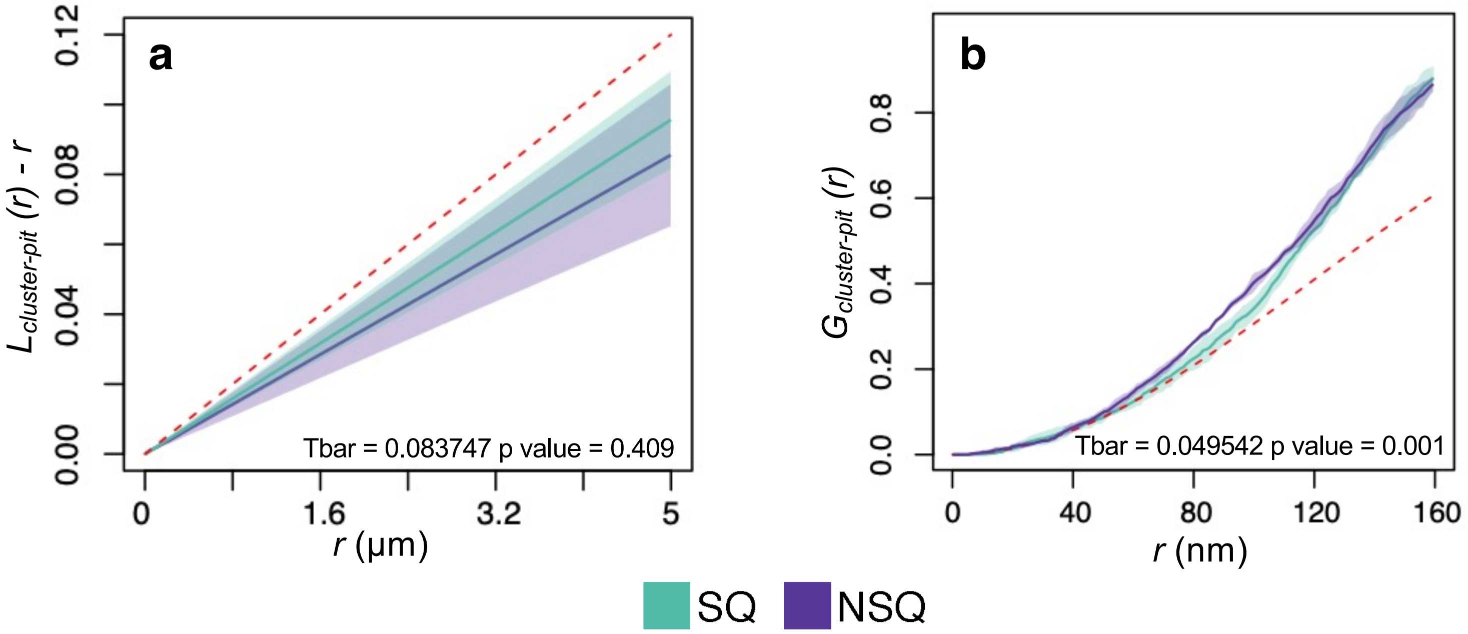


**Supplementary Figure 9.** Dependence of the spatial pattern of paxillin clusters on nanopits. Summary statistics was used to determine the dependence of the spatial patterns of paxillin clusters on nanopits. a) The $L_{cluster\text{–}pit}$ *(r)* function describes the spatial dependence of paxillin clusters on nanopits. *L(r)* > 0 indicates a positive spatial association, while $L_{cluster\text{–}pit}$ *(r)* < 0 indicates a negative spatial association. b) The $G_{cluster\text{–}pit}$ *(r)* function describes the nearest neighbour distances between paxillin clusters and nanopits. A $G_{cluster\text{–}pit}$ *(r)* value above the line representing a random and independent process denotes aggregation of paxillin clusters at the spatial scale *r*. Conversely, $G_{cluster\text{-}pit}$ *(r)* below the line representing a random and independent process denotes a regular pattern at the spatial scale *r.* The lines indicate the empirical summary function estimated from the dataset, while the shaded areas represent the confidence interval. The red line denotes the expected values for independence of the spatial location of paxillin clusters from nanopits. We present the test statistic Tbar and *P* value from the permutation test of the summary statistic.

**Supplementary Note 2**

Statistics on the spatial patterns of paxillin clusters

Robust spatial statistics require the measurement of intensity (also denoted as $\lambda$), correlation and spacing to infer the process that defines a spatial pattern[3]. We presented intensity and the $J_{cluster\text{–}pit}\ (r)$ function in the main text. We followed the excellent introduction to spatial statistics given by Baddeley et al.[3], from which the equations listed below were obtained.

We consider individual paxillin clusters to be a point within a region of interest. All paxillin clusters within a region of interest were then considered to be a finite point pattern $\mathbf{X}^{cluster}$ with an intensity (count per area) of $\lambda$. The nanopits underlying these paxillin clusters were also considered as a point pattern and denoted as $\mathbf{X}^{pit}$. From these, we calculated the following summary statistics of correlation as a function of the inter-object distance $r$. Summary statistics that describe or test the randomness of paxillin cluster positioning are denoted as $L_{cluster\text{–}cluster}\ (r)$, $G_{cluster\text{–}cluster}\ (r)$ and $J_{cluster\text{–}cluster}\ (r)$. Conversely, summary statistics that describe the spatial dependence of paxillin clusters on nanopits are denoted as $L_{cluster\text{–}pit}\ (r)$, $G_{cluster\text{–}pit}\ (r)$ and $J_{cluster\text{–}pit}\ (r)$.

The *L(r)* function

The correlation between paxillin clusters was measured through the $L_{cluster\text{–}cluster}\ (r)$ function corrected for inhomogeneous cluster intensity. The *L(r)* function is a version of the Ripley's *K* function transformed to stabilize variance and for ease of visualization. In essence, the *K(r)* function describes the expected number of neighbours of a paxillin cluster within a distance ≤ $r$.

The $K_{cluster\text{–}cluster}\ (r)$ function describes the average number of neighbours lying within a distance $r$ of a typical paxillin cluster $x$ found at location $u$:

$$K_{cluster-cluster}(r) = \mathbb{E}\left[\sum_{x \in \mathbf{X}^{cluster}} \frac{1}{\lambda(x)} \mathbf{1}\left\{0 < \left\| u - x_j \right\| \le r\right\} \mid u \in \mathbf{X}^{cluster}\right].$$

Moreover, the cross-type function $K_{cluster\text{–}pit}\ (r)$ describes the average number of nanopits lying within a distance $r$ of the typical paxillin cluster $x$ found at location $u$. $K_{cluster\text{–}pit}\ (r)$ measures the dependence of paxillin cluster location on nanopit location:

$$K_{cluster-pit}(r) = \frac{1}{\lambda_{pit}} \mathbb{E}\left[\sum_{x \in \mathbf{X}^{pit}} \frac{\mathbf{1}\{0 < \|x - u\| \le r\}}{\lambda_{pit}(x)} \mid u \in \mathbf{X}^{cluster}\right].$$

Both $K_{cluster\text{–}cluster}\ (r)$ and $K_{cluster\text{–}pit}\ (r)$ are linearized in the following manner, to obtain the *L(r)* function.

$$L(r) = \sqrt{\frac{K(r)}{\pi}}.$$

Here, we plotted *L(r) – r* for ease of visualization. *L(r) – r* = 0 indicates a lack of correlation or shows independence between objects (i.e., points generated by a random point process). *L(r) – r* > 0 indicates spatial correlation or positive association with more objects lying within the distance *r* of a typical paxillin cluster than would be expected in a random point process. Conversely, *L(r) – r* < 0 indicates spatial regularity or negative association with fewer objects lying within the distance *r* than expected in a random point process.

The *G(r)* function

Measurement of paxillin cluster spacing was performed using the *G(r)* and *J(r)* functions corrected for inhomogeneous cluster intensity. The *G* function describes the distribution of nearest-neighbour distances or the shortest distances between pairs of objects.

The $G_{cluster\text{–}cluster}$ *(r)* function is a cumulative distribution that denotes the probability of obtaining a distance ≤ $r$ between a typical paxillin cluster $x$ at an arbitrary point $u$ and its nearest neighbouring paxillin cluster.

The $G_{cluster\text{–}cluster}$ *(r)* function is calculated as follows:

$$G_{cluster-cluster}(r) = 1 - \mathbb{E}\left[\prod_{x \epsilon \mathbf{X}^{cluster} \cap b(u,r)} \left(1 - \frac{\lambda_{min}}{\lambda(x)}\right) \mid \mathbf{X}^{cluster} \text{ has a point at } u\right],$$

where $b(u, r)$ defines the disc of radius $r$ centered at an arbitrary location $u$.

Analogously, $G_{cluster\text{–}pit}$ *(r)* denotes the probability of obtaining a distance ≤ $r$ between a typical paxillin cluster $x$ at an arbitrary point $u$ and the closest nanopit $x^{pit}$ is and is defined as follows:

$$G_{cluster-pit}(r) = \mathbb{P}\{d(u, \mathbf{X}^{pit}) \leq r \mid \mathbf{X}^{cluster} \text{has a point at } u\},$$
$$d(u, \mathbf{X}^{pit}) = \min\{\left|\left|u - x^{pit}\right|\right| : x^{pit} \epsilon \mathbf{X}^{pit}\}.$$

Unlike the *L(r)* function, the *G(r)* function shows a varying line for a random point process. When *G(r)* is above the line for a random point process, it is more likely that distances between nearest neighbours are shorter than expected from a random point process. This indicates the spatial aggregation of paxillin clusters or a positive association between paxillin clusters and nanopits. Conversely, when *G(r)* is below the line for a random point process, there is a higher chance that the distances between nearest neighbours are longer than expected from a random point process. This indicates the spatial regularity of paxillin clusters or a negative association between paxillin clusters and nanopits.

The *J(r)* function

The $J_{cluster-cluster}(r)$ function deviates from the $J_{cluster-pit}(r)$ function described in the main text in the following manner. $J_{cluster-cluster}(r)$ is a ratio of the probability that the distance between a paxillin cluster and the nearest cluster is greater than *r,* and the chance of finding a cluster within a distance *r* from any fixed location $u$. $J_{cluster-cluster}(r) > 1$ denotes a regular spatial pattern of paxillin clusters, $J_{cluster-cluster}(r) < 1$ denotes a clustered spatial pattern of paxillin clusters and $J_{cluster-cluster}(r) = 1$ denotes a random pattern of paxillin cluster positioning.

The $J_{cluster-cluster}(r)$ function is defined as follows:

$$J_{cluster-cluster}(r) = \frac{1 - G_{cluster-cluster}(r)}{1 - F_{cluster-cluster}(r)},$$

where the empty space function $F_{cluster-cluster}(r)$ describes the cumulative distribution of the distances from an arbitrary location $u$ to the nearest paxillin cluster $x$:

$$F_{cluster-cluster}(r) = 1 - \mathbb{E}\left[\prod_{x \in \mathbf{X}^{cluster} \cap b(u,r)} \left(1 - \frac{\lambda_{min}}{\lambda(x)}\right)\right],$$

where $b(u,r)$ defines the disc of radius $r$ centered at $u$.

Null hypothesis testing of spatial patterns

To analyse the differences in the spatial patterns of paxillin clusters between substrates, we performed a permutation test[4]. The permutation test uses random permutations of all the point patterns in the dataset to obtain the test statistic Tbar. Tbar is a measurement of the differences between group means weighted by the estimated within-group variance. Spatial statistics and measurements of statistical significance (Berman test, maximum absolute deviation test and permutation test), were implemented using the *spatstat*[5] package for R.

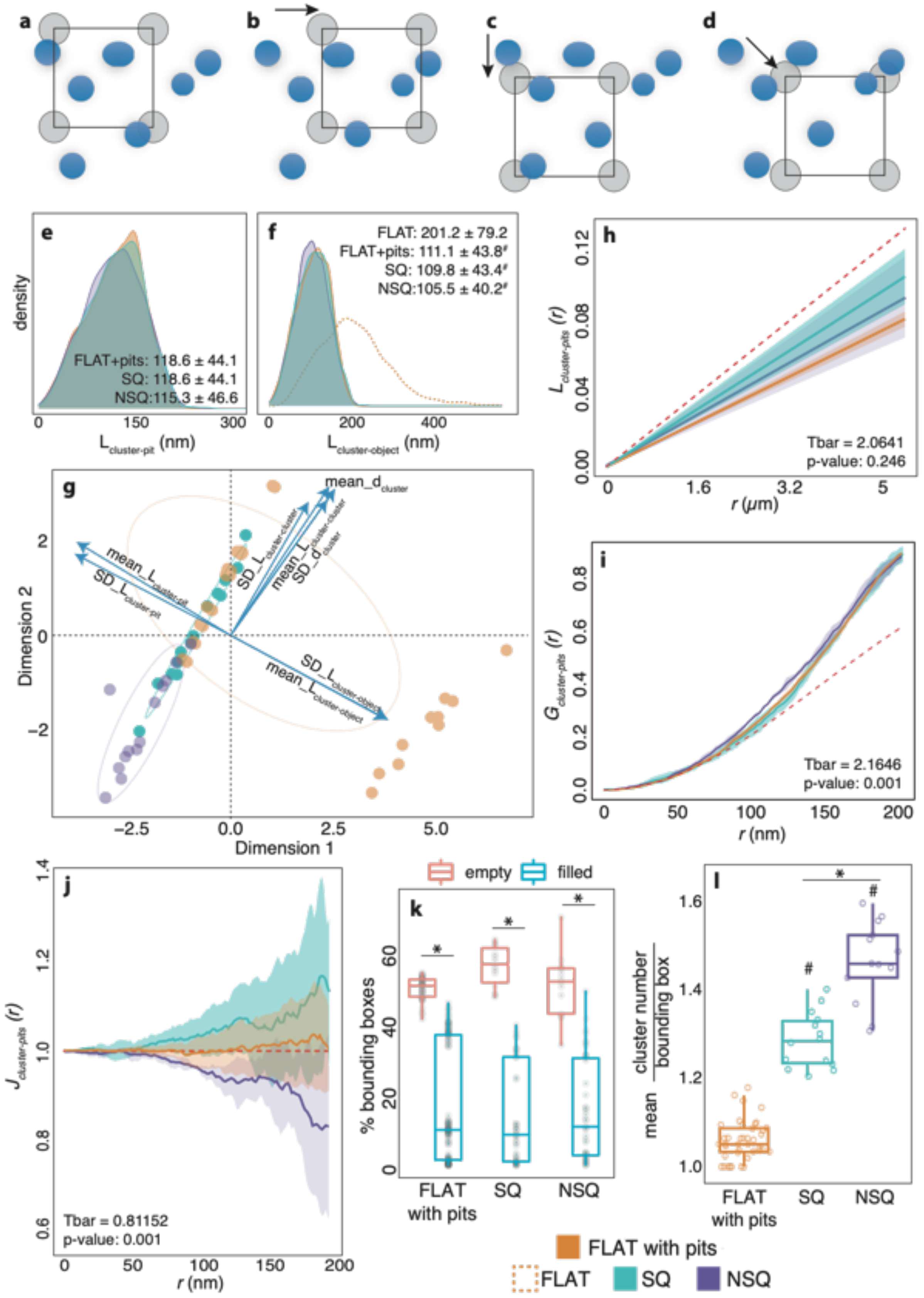


**Supplementary Figure 10.** Spatial dependence of the paxillin clusters found on FLAT on nanopits. a) To determine the importance of the nanopits on the spatial arrangement of paxillin clusters, we overlaid the paxillin clusters identified on FLAT with nanopits in a square arrangement. b–d) The nanopit locations were shifted by 150 nm in the x, y and z directions to provide four different configurations. Paxillin clusters from FLAT overlaid with nanopits were then used for further analysis. e–f) The length between a cluster and the nearest neighbouring cluster ($L_{cluster–pit}$) or nearest neighbouring object (either a nanopit or a cluster, $L_{cluster–object}$) exhibited similarity between FLAT with pits and SQ. g) Principal component analysis showed that some configurations of FLAT with pits clustered together with SQ. h) $L_{cluster–pit}(r)$ values were statistically similar between FLAT with pits, SQ and NSQ. Statistical analysis of the $L_{cluster–pit}(r)$ function for FLAT with pits revealed a significant difference from a random point process (MAD test: *P* value = 0.025). i) $G_{cluster–pit}(r)$ values were statistically significant between FLAT with pits, SQ and NSQ. Statistical analysis of the $G_{cluster–pit}(r)$ function for FLAT with pits revealed a significant difference from a random point process (MAD test: *P* value = 0.025). j) The $J_{cluster–pit}(r)$ function was statistically significant between FLAT with pits, SQ and NSQ. Statistical analysis of the $J_{cluster–pit}(r)$ function for FLAT with pits revealed statistical similarity to a random point process (MAD test: *P* value > 0.425). k) The percentage of empty and filled bounding boxes were significantly different regardless of the nanopattern. l) On average, the number of paxillin clusters on FLAT with pits was significantly lower than that on SQ and NSQ. n = 11/15/13 regions analysed for FLAT/SQ/NSQ obtained from 3 independent experiments. # Denotes statistical significance against FLAT, and * denotes statistical significance against the indicated pairs, as assessed using one-way ANOVA with Tukey's post hoc test.

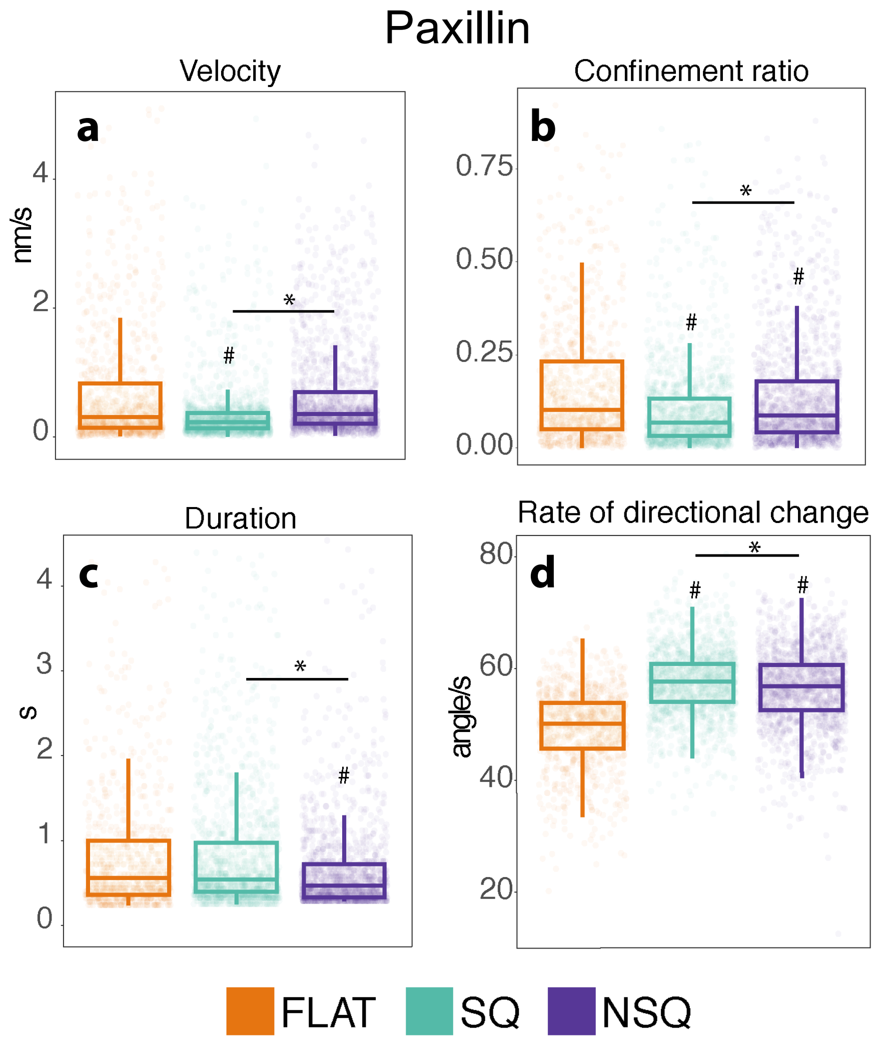


**Supplementary Figure 11.** Diffusion characteristics of paxillin. a) The velocity (nm/s), b) confinement ratio, c) duration (s) and d) rate of directional change (angle/s) of paxillin tracks were altered by nanopatterns compared with FLAT. The confinement ratio is defined as the displacement normalized to the total path length. The points denote individual paxillin tracks. The data were obtained from one independent experiment with n = 864/1534/1409 tracks from FLAT/SQ/NSQ. # Denotes statistical significance against FLAT, and * denotes statistical significance against the indicated pairs, as assessed using one-way ANOVA with Tukey's post hoc test.

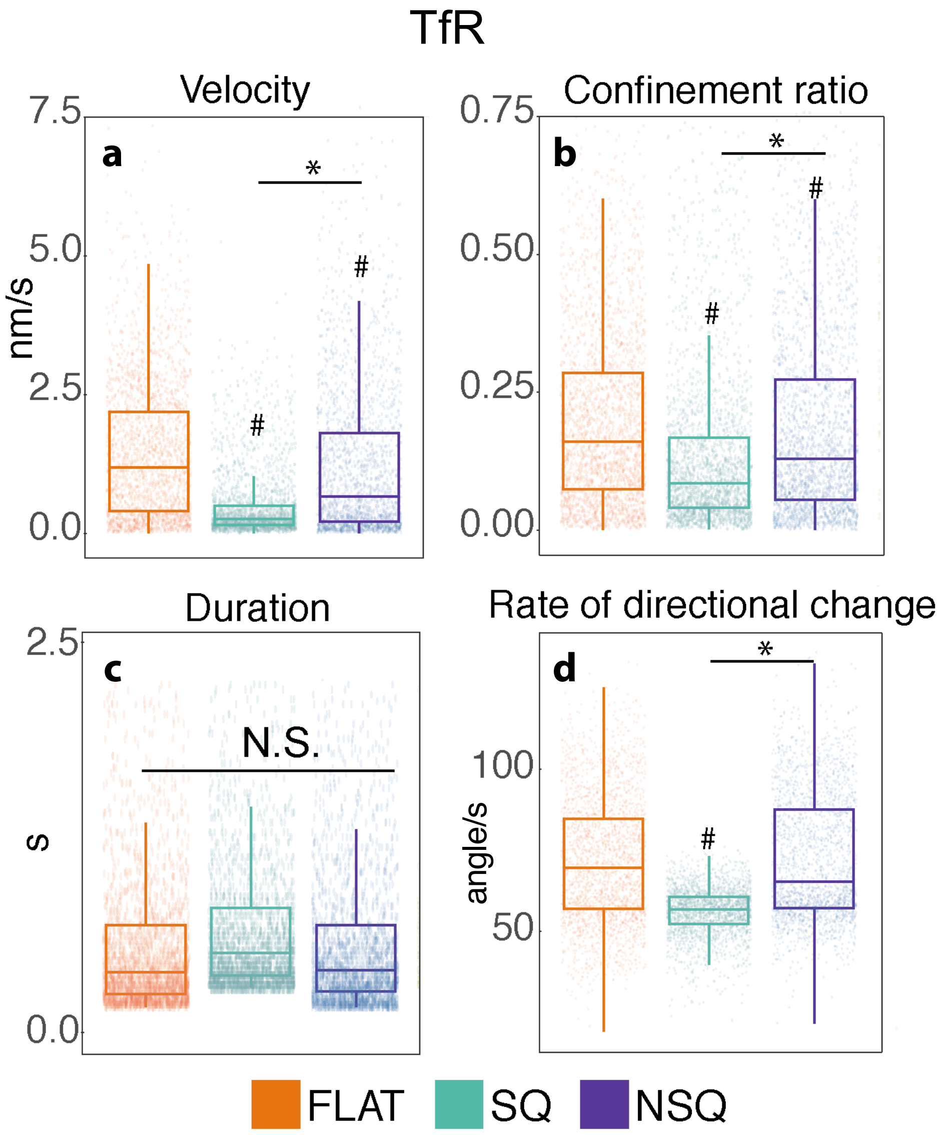


**Supplementary Figure 12.** Diffusion characteristics of paxillin. a) The velocity (nm/s), b) confinement ratio, c) duration (s), and d) rate of directional change (angle/s) of paxillin tracks were altered by nanopatterns compared with FLAT. The confinement ratio is defined as the displacement normalized to the total path length. The points denote individual paxillin tracks. The data wee obtained from one independent experiment with n = 2002/1031/1309 tracks from FLAT/SQ/NSQ. # denotes statistical significance against FLAT, and * denotes statistical significance against the indicated pairs, as assessed using one-way ANOVA with Tukey's post hoc test.

**Supplementary Table 3.** Diffusion subspecies of paxillin. Diffusion subspecies were calculated using the variational Bayesian approach for classifying single-particle trajectories.[6]

| **Topography** | **Diffusion Coefficient (um$^2$/s)** | **Occupancy (proportion of tracks)** |
|---|---|---|
| FLAT | 0.00892 | 0.48 |
| | 0.0303 | 0.341 |
| | 0.0768 | 0.119 |
| | 0.254 | 0.0465 |
| | 0.445 | 0.0144 |
| SQ | 0.0118 | 0.34 |
| | 0.032 | 0.53 |
| | 0.0796 | 0.12 |
| | 0.438 | 0.00962 |
| NSQ | 0.0178 | 0.417 |
| | 0.0503 | 0.412 |
| | 0.0793 | 0.11 |
| | 0.368 | 0.0607 |

**Supplementary Table 4.** Diffusion subspecies of transferrin receptor (TfR). Diffusion subspecies were calculated using the variational Bayesian approach for classifying single-particle trajectories.[6]

| Topography | Diffusion Coefficient ($\mu m^2/s$) | Occupancy (proportion of tracks) |
|---|---|---|
| FLAT | 0.0325 | 0.29 |
| | 0.0905 | 0.285 |
| | 0.181 | 0.174 |
| | 0.314 | 0.229 |
| | 0.473 | 0.022 |
| SQ | 0.0339 | 0.689 |
| | 0.108 | 0.253 |
| | 0.396 | 0.0576 |
| NSQ | 0.0338 | 0.51 |
| | 0.0939 | 0.378 |
| | 0.261 | 0.0862 |
| | 0.463 | 0.0252 |

**Supplementary Table 5.** Sequences of the primers used for the quantitative measurement of gene expression.

| Gene | Primer sequence | | Amplicon size |
|---|---|---|---|
| 18S ribosomal RNA (reference gene) | Fwd | AAGTCCCTGCCCTTTGTACACA | 100 |
| | Rev | GATCCGAGGGCCTCACTAAAC | |
| *BGLAP* (osteocalcin) | Fwd | CTGACCTCACAGATGCCAAG | 98 |
| | Rev | GTAGCGCCGGAGTCTGTTC | |
| *SPP1* (osteopontin) | Fwd | TCAGGACAACAACGGAAAGGG | 139 |
| | Rev | GGAACTTGCTTGACTATCGATCAC | |
| *SP7* (osterix) | Fwd | GGTCCAGGCAACACACCTAC | 184 |
| | Rev | GGTAGGGAGCTGGGTTAAGG | |
| *ALPL* (alkaline phosphatase) | Fwd | GTGCCAGAGAAAGAGAGAGA | 72 |
| | Rev | TTTCAGGGCATTTTTCAAGGT | |

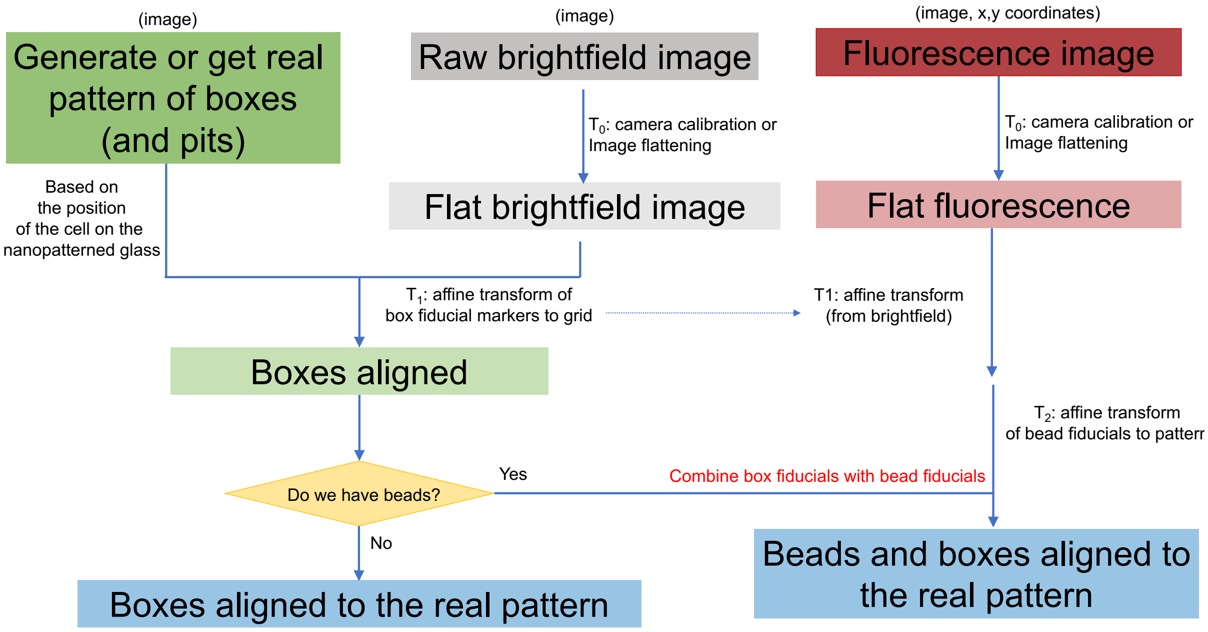


**Supplementary Figure 13.** Schematic representation of the process used for aligning the images obtained from super-resolution microscopy or single-particle tracking to the underlying nanopit substrates.

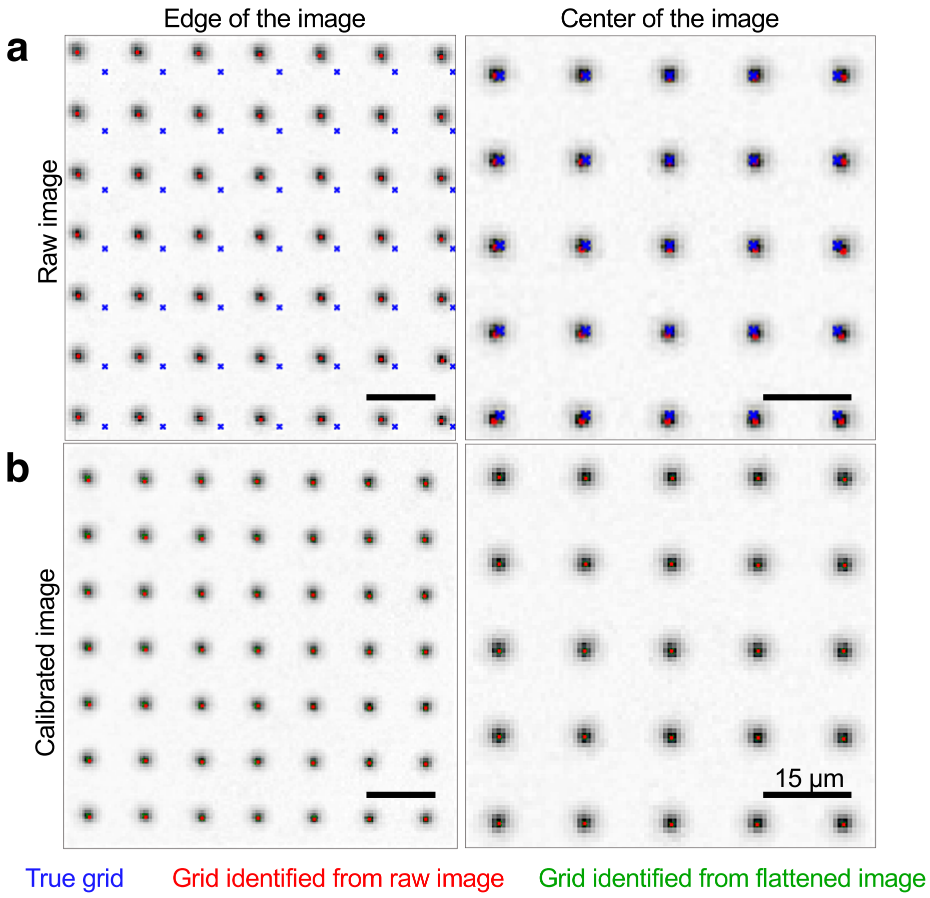


**Supplementary Figure 14.** Camera calibration to flatten images. a) A grid of regularly arrayed structures (600 nm in size) was used to calibrate the camera by calculating the transformation matrix that was required to transform the image to the true grid. b) The corrected image results in alignment of the image coordinates with the true coordinates of the grid. Calibration of the images reduced the average error between the image and true grid from 308 nm to 78 nm. The error was averaged from the distance between structures in the image and the true grid.

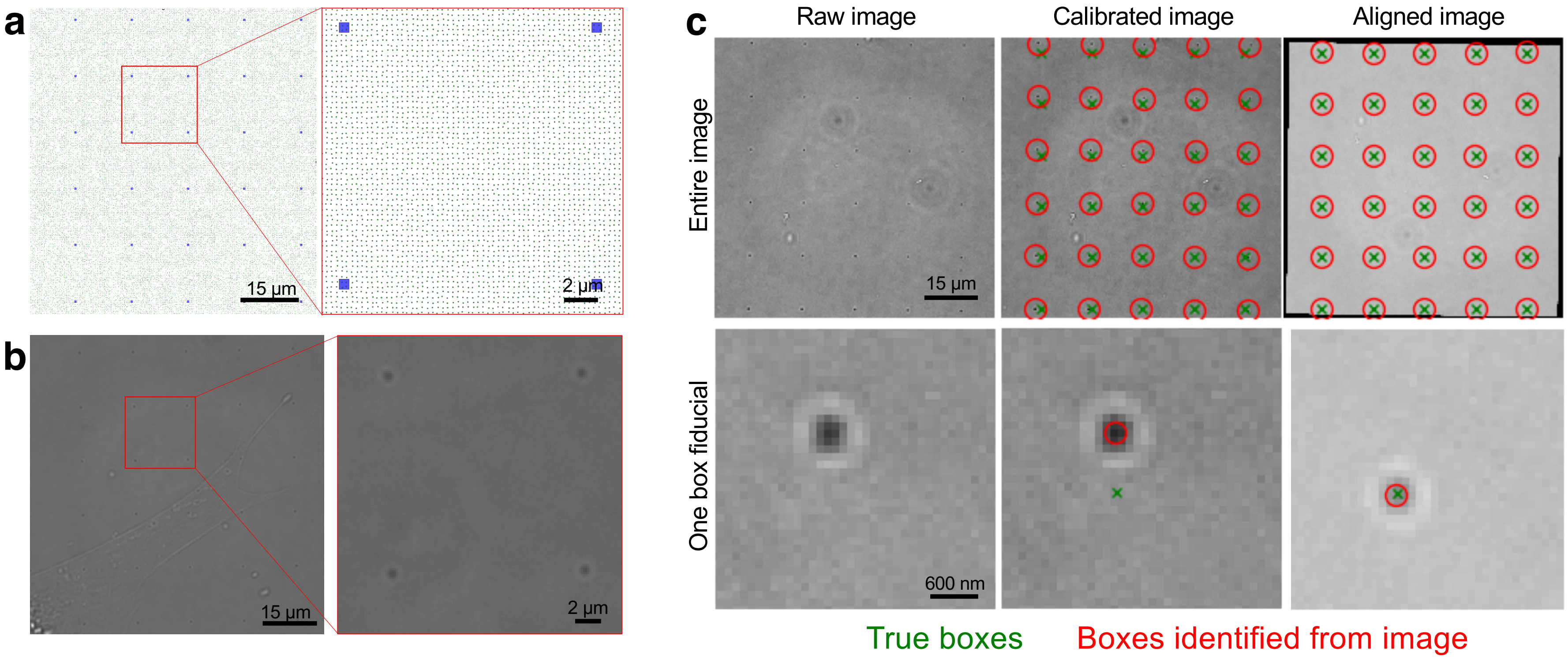


**Supplementary Figure 15.** Alignment of box fiducial markers from the brightfield image to the true nanopit pattern. a–b) Fiducial markers 600 nm in size were digitally designed and fabricated alongside the nanopits. c) After camera calibration of the bright-field image, the affine transformation that converts the calibrated image to the pattern of the true box grid was calculated. The calculated affine transformation was then applied to super-resolution images. The calibration and alignment of the images reduced the mean error of alignment between the image and true grid from 4573 nm to 74 nm. The error was averaged from the distance between the box fiducials identified in the image and the true box fiducial grid.

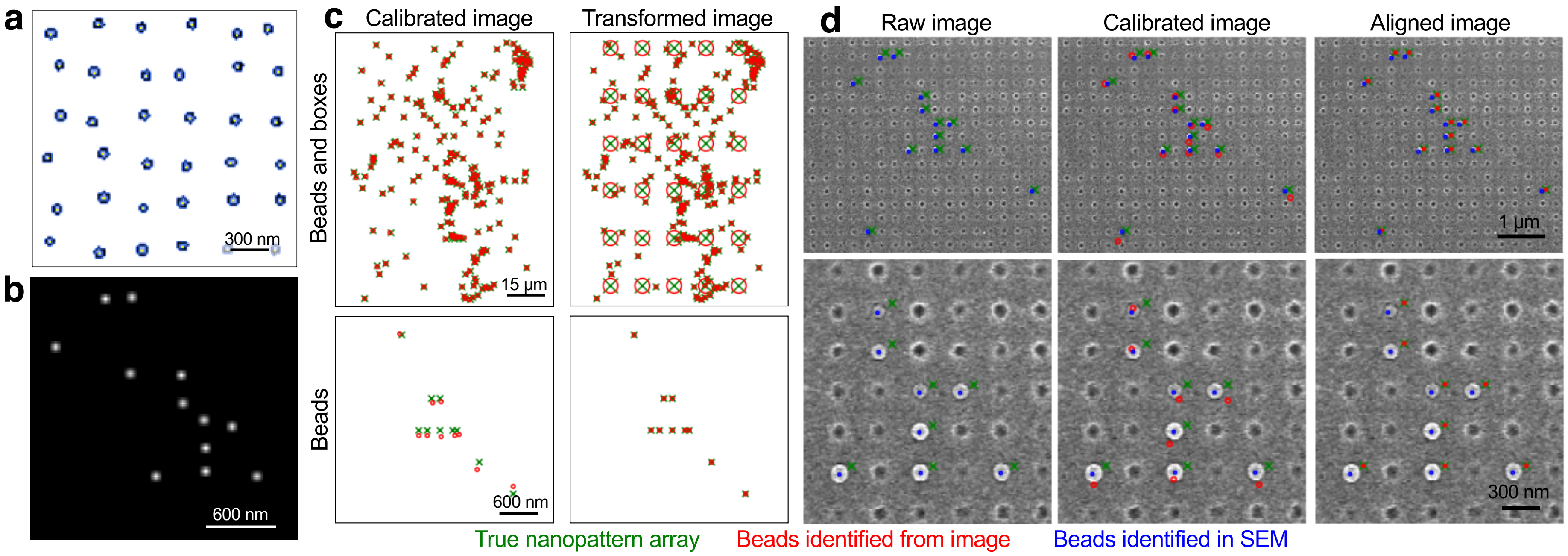


**Supplementary Figure 16.** Alignment of bead fiducial markers from fluorescence images to the true nanopattern array. a) A representative image of the digital design file containing nanopits on NSQ. b) Representative fluorescence images of the polystyrene beads immobilized in nanopits. c) Polystyrene microspheres were used as additional fiducial markers in images used in single-particle tracking. c) After camera calibration and alignment of box fiducials, the affine transformation that converts the calibrated image of the bead pattern to the true nanopattern was calculated using a random sample consensus (RANSAC) optimization method. d) Scanning electron microscopy (SEM) was used to visualize both the nanopits and the beads, thus permitting validation of the alignment method between the image and the true grid. The calibration of the images reduced the mean error of alignment between the image and true grid from 1130 nm to 70 nm. The error was averaged from the distance between the bead fiducials identified in the image and the true nanopit array.

**Supplementary Note 3**

Alignment of images to nanopatterns

Critical to the analysis performed in this study is the accurate alignment of images (from super-resolution microscopy and single-particle tracking) to the underlying patterns of nanopits. All images were first calibrated to correct the distortion of the image caused by the physical interaction of light with the microscope, the camera and the sensor. We obtained the distortion and camera matrix by comparing a bright-field image and the known coordinates of a substrate with structures with a length of 600 nm placed in a regular array with a 15 μm pitch (Supplementary Figure 14). The resulting distortion and camera matrix were applied to images captured by super-resolution and single-particle tracking prior to alignment. Camera calibration was performed using the *opencv* package (v3.4.2)[7].

An image file of the nanopatterns was first obtained from the design file used for the electron beam lithographic fabrication of the nanopatterns. Coordinates in (x, y) of the centres of box fiducials and nanopits were obtained from the image file using a template-matching algorithm in the *scikit-image* package (v0.15.0)[8]. Coordinates in (x, y) of the bead centroids were obtained using the multi-emitter maximum likelihood estimator engine of the *ThunderSTORM* plugin[9] for ImageJ. Coordinates of box fiducials (length, 600 nm; 15 μm apart in a square grid) and polystyrene beads immobilized in nanopits were identified from calibrated bright-field and fluorescent images, respectively, using template matching. The detection of the box and bead centre coordinates was enhanced to sub-pixel accuracy using the *find_peaks* function in the *astropy* package (v0.6)[10].

We then compared the coordinates of the boxes or beads identified from images with the true box patterns or nanopits using robust matching via the random sample consensus (RANSAC) algorithm. The RANSAC algorithm identifies points that correspond between the image and the true pattern by weighting the distances of possible points in a small neighbourhood. The RANSAC algorithm then provides a subset of true box and nanopattern points that correspond to the boxes or beads identified from images. The transformation needed to align the two sets of points was then estimated using an affine transformation. Camera calibration and alignment reduced the mean error of alignment between the image and true fiducials. For images of paxillin captured using super-resolution microscopy, we used the box fiducials to minimize the mean error of alignment to 74 nm (Supplementary Figure 15). For images of paxillin and the transferrin receptor captured using single-particle tracking, we used both the box and bead fiducials to minimize the mean error of alignment to 70 nm (Supplementary Figure 16). The *ransac* and *AffineTransform* functions were implemented using the *scikit-image* package (v0.15.0)[8]. The entire pipeline used for image alignment was implemented using Python 3.6 through the Spyder software (v3.3.0).

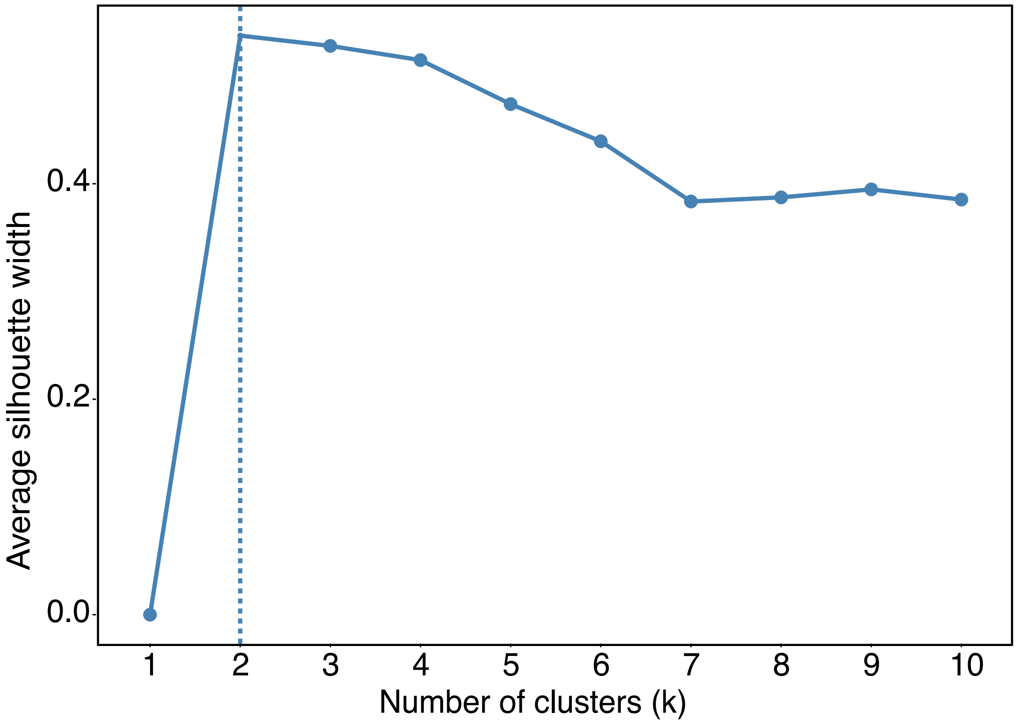


**Supplementary Figure 17.** Determination of the number of clusters (*k*) needed for *k*-means clustering. The optimum *k* was determined by the maximization of the average silhouette width and the stability of the membership of data points to a cluster.

### References


1. Püspöki, Z., Storath, M., Sage, D. & Unser, M. Transforms and Operators for Directional Bioimage Analysis: A Survey. *Adv Anat Embryol Cell Biol* **219**, 69–93 (2016).
2. Campello, R. J. G. B., Moulavi, D. & Sander, J. Density-based clustering based on hierarchical density estimates. in *Advances in knowledge discovery and data mining* (eds. Pei, J., Tseng, V. S., Cao, L., Motoda, H. & Xu, G.) 160–172 (Springer Berlin Heidelberg, 2013).
3. Baddeley, A., Rubak, E. & Turner, R. *Spatial point patterns : methodology and applications with R.* (CRC Press, Taylor & Francis Group, 2016).
4. Hahn, U. A Studentized Permutation Test for the Comparison of Spatial Point Patterns. *Journal of the American Statistical Association* **107**, 754–764 (2012).
5. Baddeley, A. & Turner, R. spatstat : An R Package for Analyzing Spatial Point Patterns. *J. Stat. Soft.* **12**, (2005).
6. Persson, F., Lindén, M., Unoson, C. & Elf, J. Extracting intracellular diffusive states and transition rates from single-molecule tracking data. *Nature Methods* **10**, 265–269 (2013).
7. Bradski, G. The OpenCV Library. *Dr Dobb's J. Software Tools* 120–125.
8. van der Walt, S. *et al.* scikit-image: image processing in Python. *PeerJ* **2**, e453 (2014).
9. Ovesný, M., Křížek, P., Borkovec, J., Švindrych, Z. & Hagen, G. M. ThunderSTORM: a comprehensive ImageJ plug-in for PALM and STORM data analysis and super-resolution imaging. *Bioinformatics* **30**, 2389–2390 (2014).
10. Price-Whelan, A. M. *et al.* The Astropy Project: Building an Open-science Project and Status of the v2.0 Core Package. *The Astronomical Journal* **156**, 123 (2018).
